# The complex octoploid Craterostigma genome and tissue-specific mechanisms underlying desiccation tolerance

**DOI:** 10.1101/2022.05.31.494158

**Authors:** Robert VanBuren, Ching Man Wai, Valentino Giarola, Milan Župunski, Jeremy Pardo, Michael Kalinowski, Guido Grossmann, Dorothea Bartels

## Abstract

Resurrection plants can survive prolonged anhydrobiosis, and desiccation tolerance has evolved recurrently across land plants as a common adaptation for survival in regions with seasonal drying. *Craterostigma plantagineum* was among the first model resurrection plants, and many of the genetic mechanisms underlying desiccation tolerance were discovered in this important system. Here, we analyzed the complex, octoploid Craterostigma (*C. plantagineum*) genome and surveyed spatial and temporal expression dynamics to identify genetic elements underlying desiccation tolerance. Homeologous genes within the Craterostigma genome have divergent expression profiles, suggesting the subgenomes contribute differently to desiccation tolerance traits. The Craterostigma genome contains almost 200 tandemly duplicated early light induced proteins (ELIPs), a hallmark trait of desiccation tolerance, with massive upregulation under water deficit. We identified a core network of desiccation responsive genes across all tissues but observed almost entirely unique expression dynamics in each tissue during recovery. Roots and leaves have differential responses related to light and photoprotection, autophagy, and nutrient transport, reflecting their divergent functions. Our findings highlight a universal set of likely ancestral desiccation tolerance mechanisms to protect cellular macromolecules under anhydrobiosis, with secondary adaptations related to tissue function.

## Introduction

Water deficit has been a pervasive challenge throughout the evolutionary history of plants, and numerous complex strategies have evolved to tolerate, escape, or avoid drought. At the extreme, resurrection plants can withstand prolonged anhydrobiosis in environments with seasonal or periodic drying such as rocky outcrops or inselbergs in Africa, Asia, Australia, and South America (1). Desiccation tolerance is an ancestral trait that likely evolved during terrestrialization (2, 3). Elements of this response have been maintained in the seeds and pollen of most flowering plants and tolerance mechanisms form the basis of core drought responses in desiccation sensitive species (4). Desiccation tolerance is widespread in non-seed plants, and it has been described in ∼78 bryophyte, fern, and fern ally families, but tolerance is relatively rare in flowering plants, arising in only nine angiosperm families (5). Over the past 30 years, more than a dozen phylogenetically diverse resurrection plants have emerged as models to study the evolution of desiccation tolerance. Since then, numerous genomic, physiologic, biochemical, and molecular studies have provided foundational insights into the complex mechanisms underlying this trait (6, 7).

*Craterostigma plantagineum* (hereon referred to as Craterostigma) was one of the earliest model resurrection plants, and it was developed by Dorothea Bartels and colleagues in the early 1990s (8). Since then, most of the genes and regulatory elements underlying desiccation tolerance in plants were discovered using molecular approaches in Craterostigma (9–13). Early discoveries were enabled by a robust and reproducible desiccation response, an efficient transformation system (14), a library of T-DNA mutants (11), and rapid screens using desiccation tolerance callus tissue (8). An early cDNA library (15) aided in molecular cloning, but the complex polyploidy of Craterostigma (2n=8x=144) has hindered development of genomic resources compared to other resurrection plant lineages (16).

Desiccation puts tremendous stress on all of the cellular macromolecules, but the stress of anhydrobiosis is not experienced uniformly across a plant. Individual tissues and cell types face unique physiological, morphological, and mechanical constraints across development and time. Light stress is the most distinguishing factor across tissues, and resurrection plants have evolved two distinct strategies to mitigate photooxidative damage. Craterostigma belongs to the group of homoiochlorophyllous resurrection plants that retain chlorophyll and thylakoid structures during dehydration. Craterostigma deploys complex molecular and biochemical mechanisms to preserve the photosynthetic apparatus while limiting the photo-oxidative damage potential of inactive chlorophyll (17, 18). This strategy enables rapid recovery upon rehydration compared to species that degrade chlorophyll and disassemble their photosynthetic apparatus (poikilochlorophylly), but it is also metabolically costly. Below ground tissues escape photooxidative stress, but they face their own unique challenges related to coordination of storage, ABA signaling, and hydraulic conductance. Most studies on desiccation and rehydration responses in resurrection plants have been conducted using leaf tissues, and responses in roots, or conserved responses across tissues are less understood.

Here, we report a high-quality assembly of the complex octoploid Craterostigma genome. Using parallel datasets in roots and leaves, we searched for shared and unique expression dynamics underlying desiccation and rehydration responses at the tissue level. We extended these analyses to the desiccation tolerant and sensitive sister taxa *Lindernia brevidens* and *L. subracemosa* respectively, to identify conserved and lineage specific responses to anhydrobiosis. Together, our results support a core desiccation response across tissues, with distinguishing differences determined by tissue functions.

## Results

### Comparative genomics of the polyploid Craterostigma genome

*Craterostigma plantagineum* is octoploid (2x=8n=112) with an estimated haploid genome size of 1.03 Gb based on flow cytometry. This translates to a compact monoploid genome size of ∼250 Mb, roughly the same size as the related diploid desiccation tolerant species *Lindernia brevidens*. We utilized a combination of PacBio long read sequencing data and HiC-based scaffolding to assemble the complex Craterostigma genome. In total, we generated 74.7 Gb or 72.5x coverage of filtered PacBio data. Raw reads were error corrected and assembled using Canu (19) with parameters optimized for haplotype assembly and phasing. The draft contigs were polished using short read Illumina data, and the resulting assembly was scaffolded using high-throughput chromatin conformation capture (Hi-C). The final assembly spans 1.71 Gb with a contig and scaffold N50 of 1.4 Mb and 5.5 Mb respectively (Supplemental Table 1). This assembly is 74% larger than the haploid genome size, indicating Canu assembled multiple haplotypes for the four homeologous regions across much of the genome. This complexity is clear in the genome assembly graph, which shows numerous interconnected edges, reflecting bubbles and ambiguities caused by haplotypes and highly similar homeologous regions (Supplemental Figure 1).

We observed numerous circular nodes in the assembly graph that correspond to complete genomes of endophytic bacteria from the Craterostigma leaf tissues. In total, we filtered out 170 Mb of microbial contamination from the assembly including 31 complete bacterial genomes, nine plasmids, and hundreds of unidentified non-plant derived contigs (Supplemental Table 2). Complete bacterial genomes have homology to well-characterized plant endophytic and rhizosphere species as well as several growth promoting species (20, 21). Craterostigma plants were propagated on soilless media and these microbiome constituents likely play important roles in plant health and possible desiccation tolerance (22).

The HiC scaffolding anchored many contigs into full length chromosomes based on synteny with *Lindernia brevidens*, but much of the genome is still in fragmented scaffolds (Supplemental Figure 2). The inability to fully scaffold the genome into 56 pseudomolecules is likely driven by chimeric read mapping and linkage in regions with high sequence homology such as homeologous chromosomes or haplotypes. The HiC scaffolded version, hereon referred to as V4 was used for annotation and downstream comparative analyses. We used the MAKER-P pipeline (23) to annotate the Craterostigma genome, with RNAseq data and protein homology as evidence. After filtering, the final annotation contains 222,749 high confidence gene models (see Materials and Methods). We assessed the annotation quality using the Embryophyta data set of Benchmarking Universal Single-Copy Orthologs (BUSCO) and found 98.4% complete proteins (1,589 out of 1,614), suggesting the annotation is high quality and largely complete.

We compared the Craterostigma genome to the closely related diploids *Lindernia brevidens* (desiccation tolerant) and *L. subracemosa* (desiccation sensitive) to explore the polyploid origin of Craterostigma and identify genomic signatures associated with desiccation tolerance within Linderneaceae. Lindernia is paraphyletic, and Craterostigma and *L. brevidens* are more closely related than the lineage containing *L. subracemosa* (24). Because of its phylogenetic position and chromosome level assembly, comparisons in Craterostigma were made primarily against *L. brevidens*. Each region of the *L. brevidens* genome has between 1-16 syntenic blocks in Craterostigma, corresponding to the octoploidy of Craterostigma and a shared whole genome duplication event in the Linderneaceae (Figure 1b). When syntenic blocks from the WGD are filtered, a maximum of eight syntenic regions in Craterostigma is observed. Only 27% of the genome is in the expected 1:4 ratio of syntenic blocks for a diploid to octoploid comparison, and 51% of the genome has between 5-8 syntenic blocks. These additional blocks correspond to assembly of multiple haplotypes for each homeologous chromosome in Craterostigma. To facilitate detailed cross-species analyses, we created a set of syntenic ortholog groups using the *L. brevidens* genome as an anchor. This approach identified 21,713 syntenic gene groupings across the three species, with 1:1 syntenic orthologs in *L. brevidens* and *L. subracemosa*, and up to 8 genes for Craterostigma. Despite differences in ploidy, the three Linderniaceae species have similar monoploid genome sizes between 250-270 Mb and repetitive element composition of 35, 34, and 31% for Craterostigma, *L. brevidens*, and *L. subracemosa* respectively. Gene density is comparable across all three species, and they maintain a high degree of gene level collinearity (Figure 1a).

**Figure 1.**
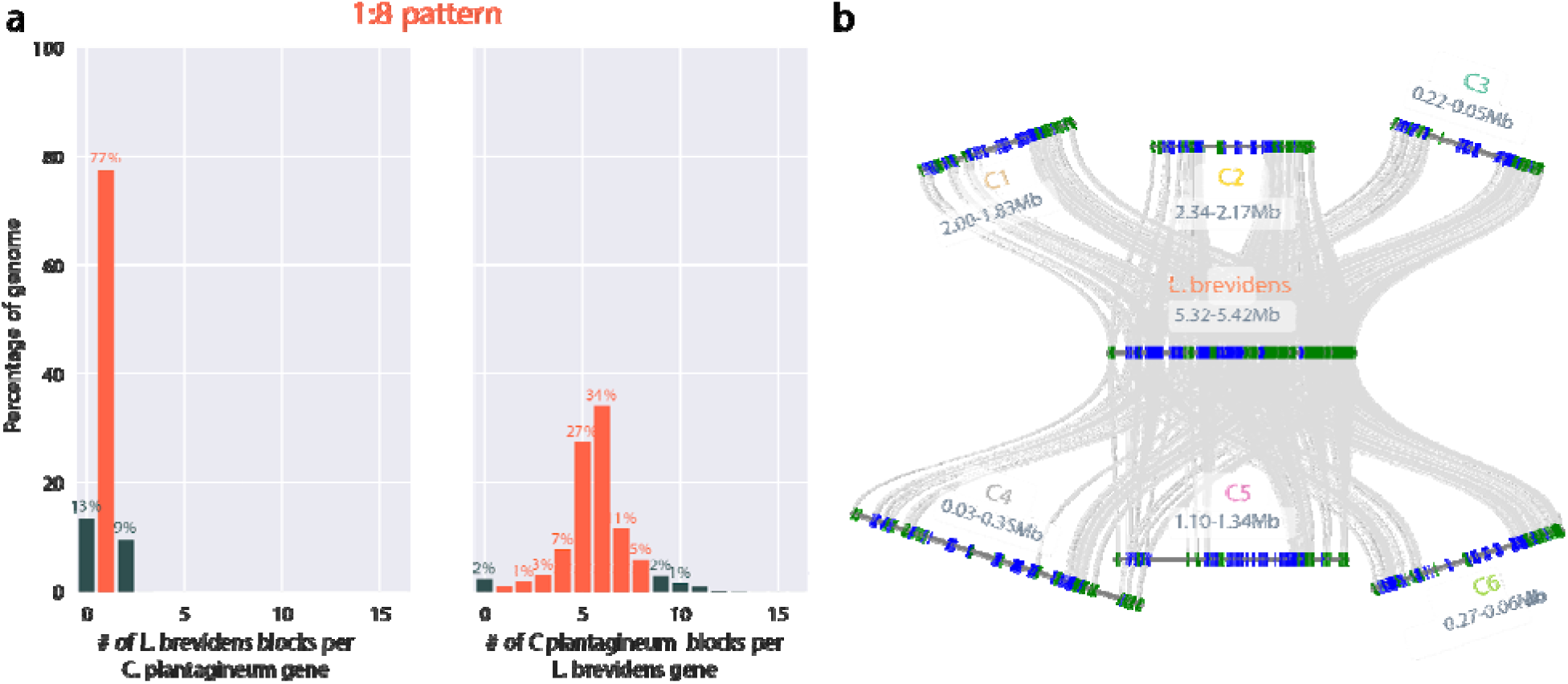
Comparative genomics and polyploidy in Craterostigma. (a) Syntenic depth comparisons between Craterostigma and *L. brevidens*. The number of *L. brevidens* blocks per Craterostigma gene is plotted on the left and the number of Craterostigma blocks per *L. brevidens* gene is shown on the right. (b) Microsynteny between one region of *L. brevidens* and six syntenic regions in Craterostigma corresponding to six of the eight haplotypes in the genome. Genes are denoted by blue and green lines (for forward and reverse orientation respectively) and syntenic gene pairs are connected by grey lines.

Many resurrection plants have polyploid genomes (16), and polyploidy is associated with adaptive potential in dynamic environments (25). We compared expression patterns of homeologous genes across the eight Craterostigma haplotypes to identify signatures of sub or neofunctionalization related to desiccation tolerance. Homeologous genes have dramatically divergent expression patterns across the surveyed tissues and desiccation/rehydration timepoints (Figure 2). Only 21% (4,500) of syntenic homeologs have identical expression patterns, with the rest showing homeolog expression bias in at least one tissue (Figure 2a). This pattern is consistent across regions ranging from 4-8 homeologs/haplotypes, but regions with eight assembled haplotypes tend to have more divergent expression profiles (Figure 2a). Homeolog expression bias varies in a tissue specific manner and is most significant in desiccated tissues (Figure 2b). Roughly one third of syntenic homeologs have distinct expression patterns in desiccated leaf or root tissues, where at least one, but not all homeologs are differentially expressed compared to well-watered tissues. This suggests that some duplicated genes have partitioned their ancestral function or have evolved new roles related to desiccation and other processes.

**Figure 2.**
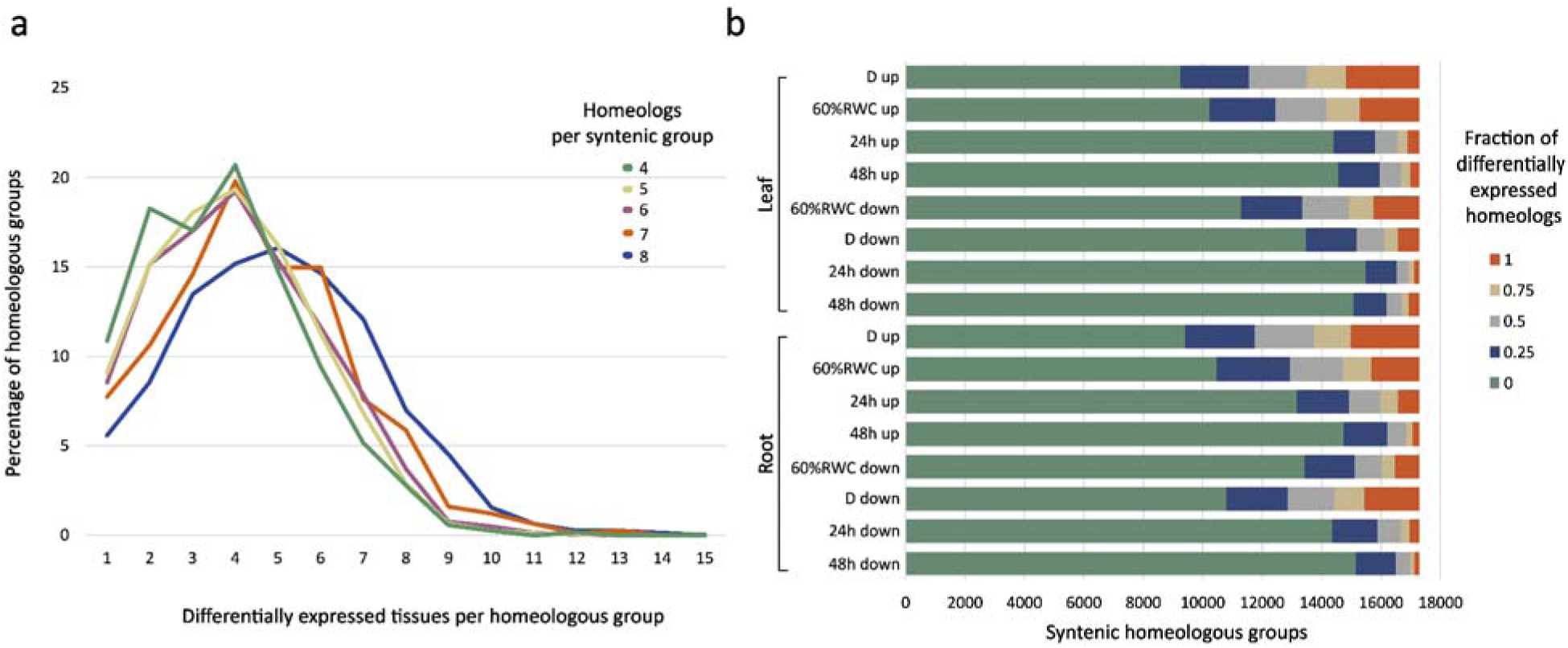
Homeolog specific expression patterns in Craterostigma. (a) The number of tissues where homeologs show different expression is plotted separately for homeologs/haplotypes ranging from a syntenic depth of 4-8. (b) Proportion of genes showing homeolog expression divergence in each of the tissue/timepoint comparisons. Syntenic groups where none of the homeologs are differentially expressed compared to well-watered are shown in green and syntenic groups where one or more genes have differential expression are shown in blue (< 25% different), grey (25-50%), and beige (50%-75). Syntenic groups showing differential expression across all homeologs compared to well-watered are shown in orange.

### Distinguishing features of desiccation response in roots and leaves

Desiccation stress presents unique physiological challenges in different organs, and plants must deploy tightly coordinated as well as tissue specific responses. Above ground tissues must mitigate photo-oxidative stress under desiccation and below ground tissues must sense changes in the regulation of water status and source-sink relations. To capture tissue specific and distinguishing responses between photosynthetic and non-photosynthetic tissues, we conducted desiccation and rehydration time course treatments for root and leaf tissues, and surveyed expression dynamics. Tissues were collected for well-watered, moderate water deficit (∼60% relative water content, RWC), desiccated (∼2% RWC), 24 hours, and 48 hours post rehydration for roots and leaves of the same plants. Leaf RWC returns to pre-stress levels within 24 hours post rehydration and plants typically resume normal photosynthetic processes within 48 hours. Three replicates were collected for each timepoint and differential gene expression and network level analyses were conducted using the RNAseq data (Figure 3a).

**Figure 3.**
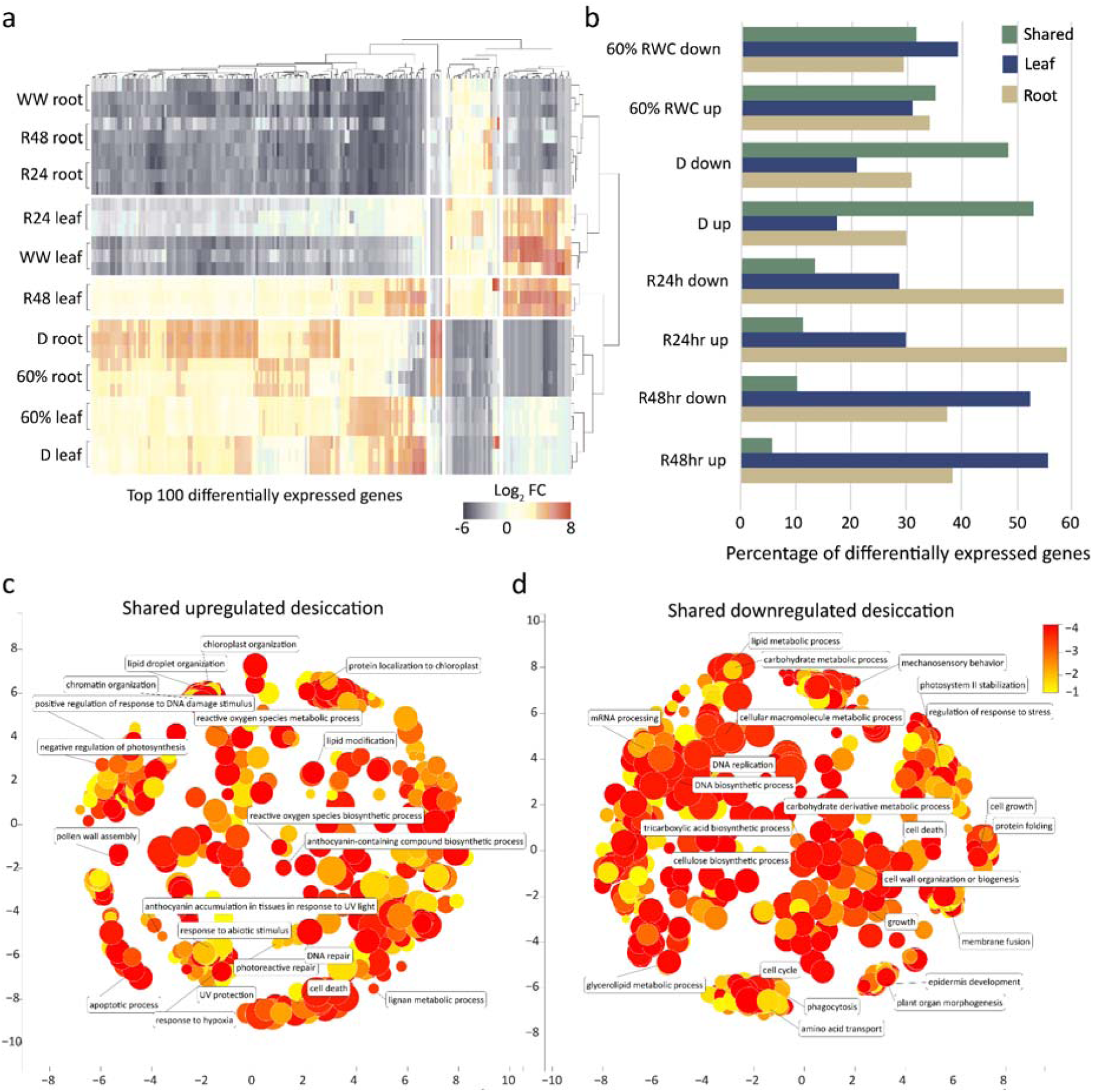
Unique and conserved expression patterns in roots and leaves during desiccation and rehydration. (a) Heatmap of the top 100 differentially expressed genes in the root and leaf timecourses. The three replicates of each sample are shown, and samples are clustered by expression patterns. (b) Overlap of shared and unique differentially expressed genes for each of the desiccation or rehydration timepoints in roots and leaves. Differentially expressed genes that are shared between roots and leaves are shown in green whereas uniquely expressed genes in leaves and roots are shown in blue and beige respectively. Enriched GO terms for shared upregulated genes under desiccation (c) and shared downregulated genes under desiccation (d). GO terms are transformed using Multidimensional Scaling to reduce dimensionality and terms are grouped by semantic similarities. GO terms with previously characterized roles in desiccation responses are highlighted. The color and size of the circles represent significance, where red represents the most significantly enriched GO terms and size correlates with the number of genes under each GO term. Clustered processes of interest are highlighted, and the full list of GO terms used to generate these graphs can be found in Supplemental Table 3.

The induction of desiccation tolerance requires the coordinated activation of hundreds of complex molecular, biochemical, and physiological responses. Consistent with this, we identified massive transcriptional reprogramming under desiccation, with 54,525 upregulated and 30,277 downregulated genes in desiccated leaves and/or roots compared to well-watered, collectively representing ∼38% of genes in the Craterostigma genome (Figure 3). Similarly high levels of differential expression were observed under moderate drought stress (60% RWC; 17,395 and 35,490 up and downregulated respectively) with comparatively fewer differentially expressed genes during rehydration in leaves and roots. We further classified differentially expressed genes as shared or unique between roots and leaves to identify conserved desiccation responses across the plant and tissue specific responses (Figure 3a and 3b). Desiccated tissues have a strong overlap of differentially expressed genes, with 53 and 48% of genes similarly upregulated or downregulated in both tissues. 35% of genes are similarly upregulated and 32% downregulated in both leaves and roots at 60% RWC. Expression dynamics in rehydrating tissues are almost entirely different in roots and leaves with only 11 and 5% of genes similarly upregulated and 13 and 10% similarly downregulated 24 and 48 hours post rehydration.

We searched for enriched gene ontology (GO) terms to identify shared and unique molecular responses in roots and leaves under desiccation and rehydration. GO terms provide a somewhat superficial overview, so we utilized Multidimensional Scaling to reduce dimensionality and group similar terms by semantic similarities (26). Among the upregulated genes in leaves and roots are many processes previously linked to desiccation tolerance including ROS detoxification, DNA repair, lipid droplet biosynthesis, osmoprotectant biosynthesis (Figure 3c; Supplemental Table 3). Interestingly, we observed some shared GO terms related to photosynthetic processes in roots and leaves, though this overlap is dwarfed by photosynthesis related GO terms in leaf specific sets. Many downregulated processes are common between roots and leaves including biosynthesis of DNA, carbohydrates, cellulose, and amino acids, and numerous growth and developmental activities (Figure 2d). Together, this highlights a core set of desiccation-related mechanisms that function across vegetative tissues to survive anhydrobiosis.

Although desiccation timepoints show substantial overlap between roots and leaves, there are important processes that are uniquely upregulated in each tissue (Supplemental Table 4, 5). The most notable difference in leaves are responses related to photosynthesis and photoprotection. Under progressive drought and desiccation, numerous GO terms related to UV and light responses, chloroplast organization, oxidative stress, photoprotection, and photosystem II stability are only upregulated in leaves. Root specific GO terms upregulated under desiccation include responses to trehalose, stress starvation, response to abscisic acid, lipid and various transport processes.

We searched for genes annotated in significant GO terms which might help us understand the specificities underlying the processes of desiccation and rehydration in Craterostigma (Supplemental Figure 5). We focused on early responsive genes, where among many gene families the abscisic acid signaling pathway-related genes MARD1-like and PYL4-like were commonly expressed in both tissues. Similarly, phosphoinositide phosphatase SAC8 transcripts were also up-regulated during initial desiccation in both leaves and roots. In *A*.*thaliana*, SAC8 is involved in auxin signaling, embryo and seed development, as well as maintenance of PtdIns4P and PtdIns(4,5)P2 levels (27). Root specific up-regulated genes include orthologs to WRKY19, the putative interacting partner with 14-3-3 proteins responsible for developmental, defense, and stress response (28), and mitochondrial LYR motif like gene responsible for Complex I Assembly (29). Among early responsive genes, we found several up-regulated candidates including hornerin-like and ankyrin repeat-containing proteins, which are involved in abscisic acid signaling pathways, ROS production, or might act as calcium-binding proteins (30, 31). Selected down-regulated transcripts in leaves belong to Dedol-PP synthase 6 and IBH1, which are involved in protein glycosylation, or regulate the cell elongation, respectively (32, 33).

Upon rehydration in roots, we noticed a change in expression of genes involved in secondary metabolism and growth regulation (Supplemental Figure 6). During recovery, root specific up or down-regulated transcripts included cytokinin dehydrogenases (AXX17_At2g38780) which are known for growth regulatory function, negatively impacting root growth and positively shoot growth, root and shoot architecture (34, 35). Rehydration in roots led to high expression of AOP1-like orthologs involved in glucosinolates synthesis (36), caffeoylshikimate esterase-like orthologs involved in lignin biosynthesis (37), S-acyltransferases responsible for protein S-acylation (38), microtubule-associated protein WVD2-like 4 regulating cytoskeleton organization (39), eIF-3 complex transcripts involved in cell proliferation (40), and proteasome inhibitors that regulate autophagy (41). These genes show distinct expression patterns in leaves, being with high expression under the drying and desiccated states (Supplemental Figure 6). Together, this highlights the large overlap in desiccation-related mechanisms between roots and leaves and unique processes under recovery related to photosynthetic and transport processes in leaves and roots respectively.

Late Embryogenesis Abundant (LEA) proteins are small, hydrophilic polypeptides that play important roles in protein protection during seed development, various abiotic stresses, and desiccation tolerance (42, 43). We observed distinct expression patterns across the different LEA subfamilies in Craterostigma roots and leaves (Supplemental Figure 3). LEAs are generally upregulated under drought and desiccation across tissues with the exception of LEA2 subfamily members which have low expression in the surveyed timepoints. Interestingly, LEA1, LEA2, LEA4, LEA5, LEA6, seed maturation protein (SMPs), and dehydrin (DHN) subfamilies have significantly higher expression in roots than comparable leaf tissues across all hydration levels (Supplemental Figure 3). This pattern is strongest in the SMP subfamily, where the six SMP orthologs have 2-to-17-fold higher expression in roots compared to leaves under the 60% RWC and desiccated timepoints. This suggests that LEAs have a central role in desiccation tolerance at a cellular level, but their accumulation varies across tissues, and may be related to organ function.

Autophagy promotes desiccation tolerance through nutrient cycling, toxin removal, and suppression of programmed cell death (44), but this response is tissue specific in the resurrection grass *Tripogon loliiformis* (45). Trehalose induces autophagy in Tripogon leaves, but roots have no autophagy responses under desiccation, and increased resource allocation is hypothesized to be sufficient for stress tolerance (45). We observed the opposite pattern in Craterostigma, where autophagy associated genes are significantly upregulated in roots compared to leaves across hydration states (Supplemental Figure 7). We identified 510 Craterostigma orthologs of the 40 autophagy-related genes (ATGs) in Arabidopsis (46), and tracked their expression patterns. Some ATGs have constitutive expressions across the surveyed timepoints, but most are upregulated under water deficit and during recovery, with noticeable differences in leaves and roots (Supplemental Figure 7). Orthologs of ATG5 and ATG8, which play a central role in carbon and nitrogen nutrient cycling, have the highest expression under desiccation and rehydration, with up to 20-fold difference in comparable leaf and root timepoints. Related to this, senescence related genes such as STR16 are significantly downregulated in roots and leaves during desiccation.

We utilized the root and leaf datasets to construct a co-expression network and identified shared and unique modules in each tissue. Six optimized clusters were identified using clust (47) with four shared between the parallel root and leaf datasets and one specific to each tissue (Figure 4). Two of the shared clusters (C0 and C1) correspond to genes induced under drought and desiccation and the other two shared clusters (C4 and C5) have high expression in well-watered and recovered tissues but decrease under water deficit (Figure 4a). The root and leaf specific clusters (C2 and C3, respectively) have high expression in well-watered tissues but decrease during progressive drought and recovery. Enriched GO terms in the leaf specific cluster include photosynthesis and photosystem repair, flavanoid and phenylpropanoid metabolism, and oxidative stress responses among others (Figure 4b). The root specific cluster has enriched GO terms related to gravitropism, nutrient transport, abiotic stress responses (Figure 4c). This cluster also contains orthologs of the negative regulators of cell death including Bax inihibitor-1 (BI-1) (48) and defender against apoptotic death 1 (DAD1) (49), further highlighting the important role of autophagy and programmed cell death in root tissues compared to leaves.

**Figure 4.**
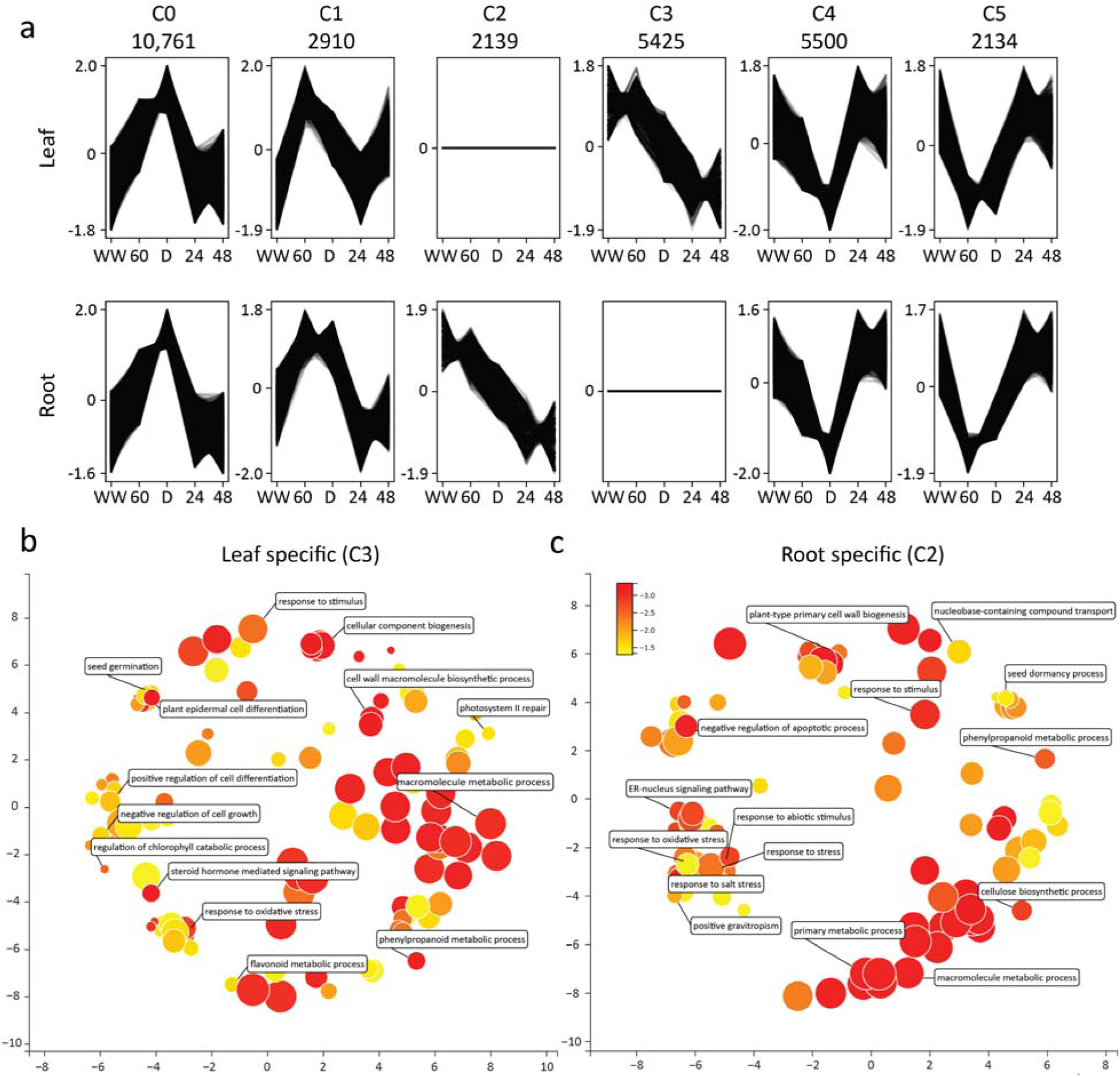
Gene Co-expression dynamics between leaf and root networks. (a) Clustering of gene expression profiles across the leaf and root desiccation and rehydration timecourses. Genes were clustered into discrete modules using clust and the transformed expression patterns are plotted for each of the 6 modules. Enriched GO terms for the leaf specific (b) and root specific modules (c). WW: well-watered; 60: moderate desiccation (∼60% relative water content); D: dehydration; 24: 24 hours post rehydration; 48: 48 hours post rehydration.

### Comparative responses to desiccation across Linderniaceae

Desiccation tolerance is an ancestral trait within some Linderniaceae lineages, and it evolved before the divergence of Craterostigma and *L. brevidens. L. brevidens* is endemic to the montane rainforests of Tanzania and Kenya where it never experiences seasonal drying but has peculiarly maintained desiccation tolerance (50). We compared the pattern of desiccation -induced gene expression across Linderniaceae to test if similar expression dynamics are observed in the two desiccation tolerant species compared to the desiccation sensitive outgroup *L. subracemosa*. We first surveyed the global expression patterns across the three species using the set of shared syntenic orthologs described above. To account for differences in ploidy, the expression values of homeologs/haplotypes in Craterostigma were averaged to achieve a single expression value for groups of 1:1:1 orthologs across the three species. This expression matrix of 21,713 syntenic orthologs was used for dimensionality reduction and network analyses. Similarly, syntenic orthologs were classified as up or downregulated if one or more Craterostigma orthologs were differentially expressed to account for sub or neofunctionalization between homeologs.

We applied a principal component analysis to the combined expression matrix and identified broad patterns of desiccation and rehydration dynamics across the three Linderniaceae species. The first two principal components collectively explain 52% of the variance and separate the expression datasets by both species and hydration level (Figure 5). Well-watered RNAseq samples form a well-defined cluster across all three species, reflecting a broad similarity of expression under normal conditions (Figure 5a, b). The 24 and 48 hour post rehydration samples from Craterostigma are also found in this cluster, suggesting many genes return to normal expression relatively quickly compared to *L. brevidens*, where recovery is slower (50, 51). The drought/desiccation and many rehydration samples are found in dispersed but distinct orthogonal clusters for each of the three species. The Craterostigma and *L. brevidens* drought/desiccation timepoints have overlapping ranges in the first principal component whereas the comparable *L. subracemosa* samples form a tight and highly distinct cluster.

**Figure 5.**
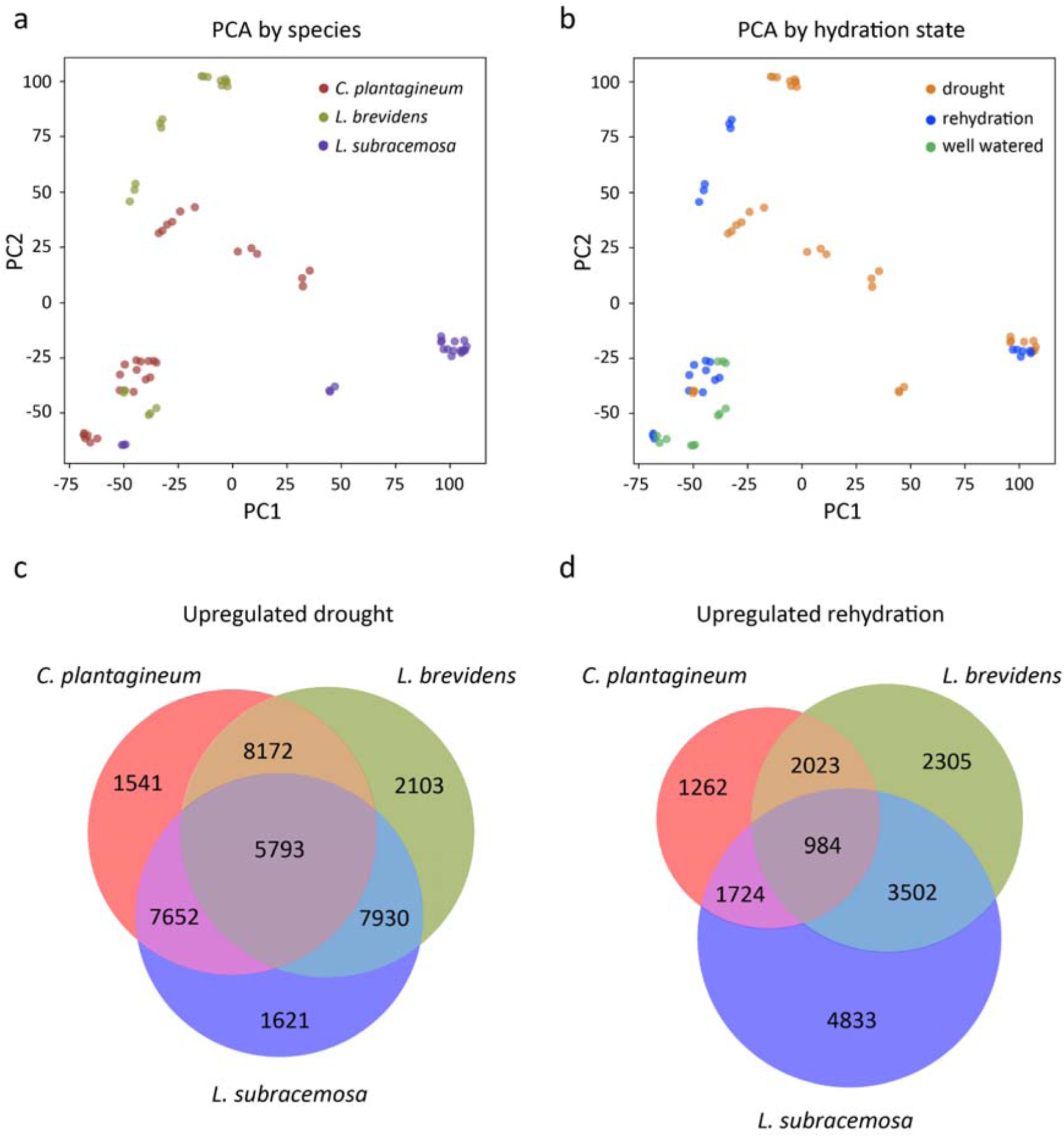
Comparative expression dynamics across the Linderniaceae under desiccation and rehydration. Principal component analysis of syntenic ortholog expression patterns colored by species (a) and water deficit (b). Expression values for syntenic orthogroups were transformed by z-score prior to principal component analysis. The first two principal components (explaining 31.0 and 21.6% of variation, respectively) are plotted for the two desiccation tolerant Lindernieaceae species (Craterostigma and *L. brevidens*) and the desiccation sensitive outgroup (*L. subracemosa*) across comparative expression datasets. Venn diagram of commonly upregulated syntenic orthogroups under drought/desiccation timepoints (c) and rehydration (d) across the three species.

These gene expression timepoints were also analyzed in a pairwise fashion to identify overlapping expression dynamics across the three species under similar hydration timepoints. We identified a core set of 5,794 orthogroups that have upregulated expression under drought across all three species regardless of desiccation tolerance or sensitivity (Figure 5c and 5d). The two desiccation tolerant species have a set of 8,172 orthologs with upregulated expression that were not found in the sensitive species (Figure 5c). Genes upregulated under rehydration show comparatively less overlap, with only 984 upregulated across all three species and 2,023 between the two tolerant species (Figure 5d). *L. subracemosa* tissues fail to recover from desiccation, and the high number of uniquely upregulated genes (4,833) may represent chaotic regulation or spurious expression in tissues that fail to recover from terminal drought stress. The generally low overlap of expression dynamics during rehydration may reflect species specific responses to repair, recovery, and resumption of normal cellular processes.

We leveraged the time ordered desiccation and rehydration datasets to construct a cross-species comparative co-expression network. This approach identified five gene clusters collectively spanning 5,497 syntenic orthogroups across the three species (Supplemental Figure 4). *L. brevidens* and Craterostigma generally have similar eigengene expression profiles across the five clusters compared to chaotic or unordered expression in *L. subracemosa*. Cluster 0 for instance, has a peak of expression under desiccation in *L. brevidens* and Craterostigma, but continually increasing expression in *L. subracemosa*. Similarly, Cluster 4 shows decreased expression under desiccation compared to well-watered and recovered in *L. brevidens* and Craterostigma but has continually decreasing expression in *L. subracemosa*. This again highlights the lack of adequate preparation to anhydrobiosis in *L. subracemosa*, and the unordered expression preceding imminent death.

### Independent duplication of ELIPs across Linderniaceae

The evolution of desiccation tolerance is associated with massive duplication of early light induced proteins (ELIPs), which play a central role in photoprotection under prolonged drying (52). All sequenced resurrection plant genomes contain large tandem arrays of ELIPs, including *L. brevidens* which has 26 ELIPs, with most found in a single tandem array of 19 copies, compared to only four in the genome of the desiccation sensitive relative *L. subracemosa*. The Craterostigma genome has 186 annotated ELIPs, and all but 24 are tandemly duplicated. Most tandem arrays are found in five haplotypes of a single homeologous region, with copy numbers ranging from nine to 18 in each haplotype (Figure 6).

**Figure 6.**
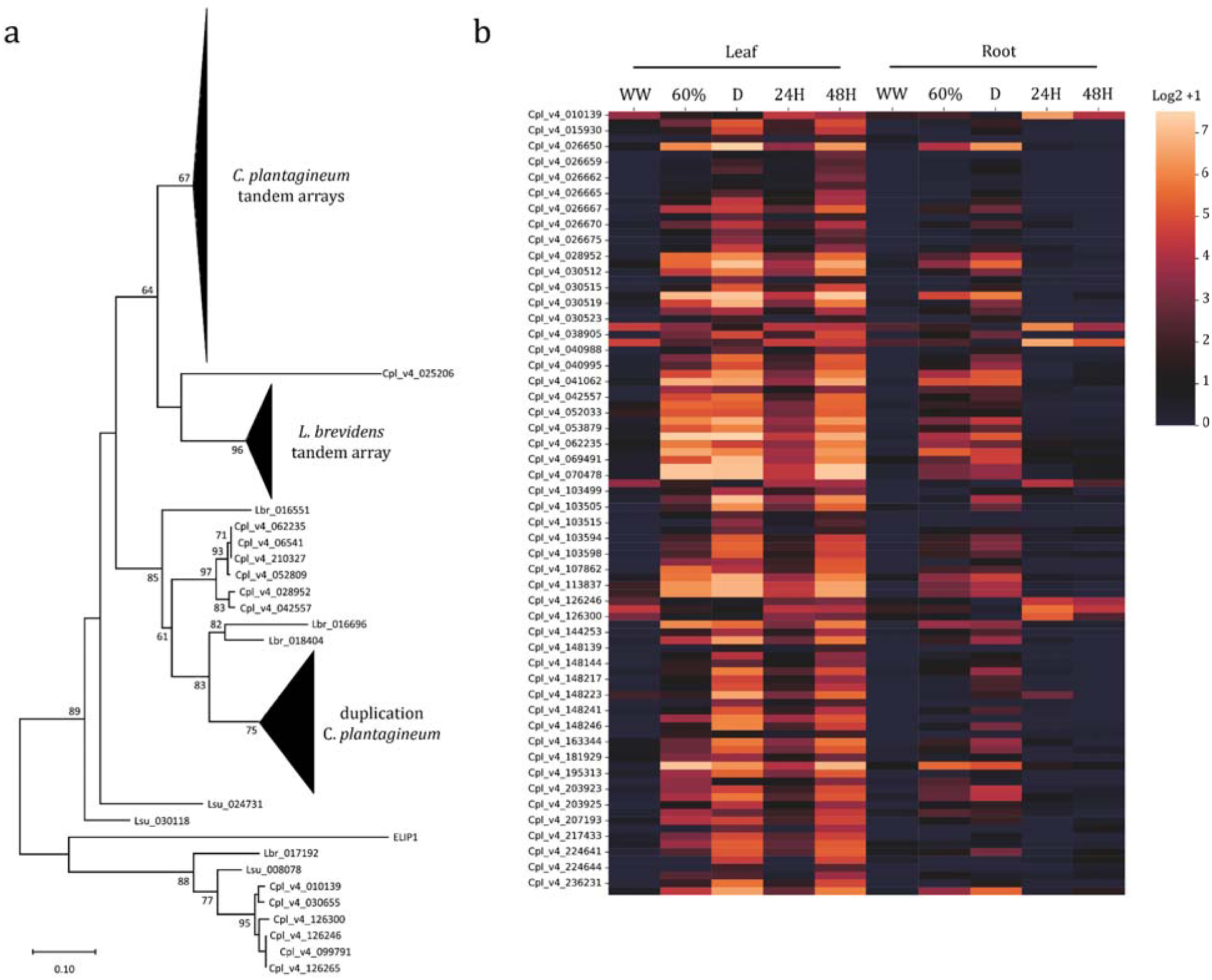
Independent tandem duplication of ELIPs in *L. brevidens* and Craterostigma. (a) Maximum likelihood phylogeny of ELIP proteins in Craterostigma, *L. brevidens*, and *L. subracemosa*. Bootstrap support values are shown for 1000 replicates. Nodes containing the large tandem arrays of ELIPs in Craterostigma and *L. brevidens* are collapsed. (b) Heatmap of Log2 transformed expression data for Craterostigma ELIPs across leaf and root desiccation and rehydration timecourses.

We surveyed the evolutionary history of ELIPs to see if the tandem duplication event is shared between *L. brevidens* and Craterostigma or if tandem duplication occurred independently. The tandem array of 19 ELIPs in *L. brevidens* is found on chromosome 8, and each of the homeologous regions in Craterostigma have only a single syntenic ELIP. The large tandem arrays of ELIPs in Craterostigma are found in a homeologous region of Chromosome 11, and *L. brevidens* has no ELIPs that are syntenic to this region. A maximum likelihood phylogenetic tree supports this finding, with a clear separation of tandemly duplicated ELIPs into distinct evolutionary clades in both species (Figure 6a). Interestingly, although the ELIP arrays have different evolutionary origins, chromosomes 8 and 11 are derived from a shared whole genome duplication event in the Linderniaceae (51). These ELIP orthologs may have novel features that make them predisposed to be duplicated during the evolution of desiccation tolerance.

We surveyed the expression of ELIPs across the leaf and root desiccation and rehydration timecourses. ELIPs have generally low or undetectable expression under well-watered conditions, but are among the most highly expressed genes during drying and recovery (Figure 6b), consistent with patterns in other resurrection plants (52). Expression of ELIPs is not uniformly high, and varies between copies within a tandem array and between arrays in homeologous regions. ELIPs have significantly lower expression in root tissues under desiccation with few being differentially expressed compared to well-watered conditions (Figure 6b). This supports the hypothesized role of ELIPs in photoprotective processes under desiccation. ELIPs are still highly expressed 48 hours post rehydration, where the tissues have returned to normal RWC, and they likely function to protect the photosynthetic apparatus as it is restored.

## Discussion

Surviving desiccation requires an intricate coordination of numerous processes and pathways to prepare tissues for prolonged anhydrobiosis. Our comparative analyses identified a broadly conserved desiccation response across tissues, highlighting the universal constraints that anhydrobiosis imposes at a cellular level. Across leaves and roots, we identified numerous pathways and processes with similar upregulation under water deficit, including many with well-defined roles in desiccation. The shift from a fluid to solid state under desiccation drastically shifts the biochemistry and mobility of cellular macromolecules. Preparation for this change requires all cell types, regardless of function, to enact a similar cascade of protective mechanisms if they are to survive anhydrobiosis. Many of the shared, desiccation-induced transcripts we identified overlap with seed maturation related processes. This includes selected LEA proteins, osmoprotectants, ROS detoxifiers, lipid body components, and the ABA-regulon among others. This finding supports the long standing hypothesis that vegetative desiccation tolerance evolved through rewiring seed pathways (2), though this has been recently contested (53). Here, we propose that a network of ancestral desiccation related mechanisms is induced in all tissues that survive anhydrobiosis (e.g. seeds, pollen, roots, leaves, etc.) as a common adaptation to protect the cellular macromolecules.

Despite the synchronization in expression between roots and leaves during desiccation, we identified almost no overlap in expression dynamics in these tissues under rehydration. While the endpoint of desiccation is the same across tissues (i.e., solid state biology), rehydration marks a return to the distinct cellular processes underlying each tissue. Many of the upregulated genes in leaves under rehydration are related to photosynthesis including photosystem repair, chlorophyll metabolism, photoprotection, and membrane repair. Root-related changes under rehydration are almost entirely distinct and include phenylpropanoid and cellulose metabolism, transport, response to gravitropism, ABA signaling, and root growth. These dynamics reflect the unique constraints that light stress and cellular function places on different tissues during rehydration.

Most molecular and multi-omics based studies in resurrection plants focus on responses to desiccation in leaves and comparatively limited work has been done in roots. Roots and leaves of the resurrection grass *Tripogon loliiformis* utilize different mechanisms to survive desiccation including contrasting induction of autophagy, sugar metabolism, and other source-sink fluxes (45). Contrasting Tripogon, we observed the upregulation of genes related to autophagy and programmed cellular recycling in both leaves and roots. This may represent distinct desiccation strategies across these highly divergent lineages, or a previously overlooked commonality.

Tandem duplication of ELIPs is a hallmark of desiccation tolerance, and the independent duplication of ELIPs in each sequenced resurrection plant supports the convergent evolution of this trait (52). Consistent with this pattern, the Craterostgima genome has massive tandem arrays of ELIPs across each of its homeologous chromosomes. Expression of ELIPs is largely confined to leaf tissue under desiccation and recovery, highlighting their role in photoprotection. ELIPs have a clear and essential role in the evolution of desiccation tolerance but whether their duplication induces desiccation tolerance or if it improves fitness of lineages with tolerance is unclear. Desiccation tolerance evolved before the divergence of Craterostigma and *Lindernia brevidens*, but duplication of ELIPs occurred after their divergence using different ancestral gene copies. This suggests that desiccation tolerance evolved in this lineage before the duplication of ELIPs, and this duplication may represent a second step for improved photoprotective capacity.

The complex polyploidy of Craterostigma has hindered genome sequencing efforts, but numerous genes, regulatory elements, and changes in the biochemistry, metabolism, and physiology underlying desiccation have been discovered in this important species. The genomic and expression datasets presented here represent key foundational resources that can link, at a systems level, genomic patterns with mechanistic findings to develop an emergent model of the pathways underlying desiccation tolerance.

## Methods

### Plant growth conditions and sampling

*Craterostigma plantagineum* plants were grown as described previously (8). Plants were propagated vegetatively and maintained under day/night temperatures of 22 and 18°C, respectively and under fluorescent lighting with an intensity of 80 μE m−2 s−1 and a photoperiod of 16/8-hr. For desiccation timecourses, plants were allowed to gradually dry in pots until reaching a partial dehydration (∼60% RWC) for the first sampling and full desiccation (D). For the rehydration timepoints, desiccated plants were allowed to rehydrate by submergence in water for 24 hours and sampled at 24 hours (R24) and 48 hours (R48) post-rehydration. Three independent biological replicates were collected for each timepoint for gene expression and relative water content datasets. Samples for RNAseq were collected at the same time each day, flash frozen in liquid nitrogen, and stored at −80°C before processing. Relative water content was determined as previously described in Bernacchia et al. (54).

### RNA extraction, library preparation, and sequencing

Total RNA was extracted from the root and leaf tissues described above using the Omega-biotek E.Z.N.A. Plant RNA Kit (Omega-biotek), according to the manufacturer’s instructions. RNA quality and quantity were assessed using gel electrophoresis and the Qubit RNA IQ Assay (ThermoFisher), respectively. RNAseq libraries were constructed using two micrograms of total RNA with the Illumina TruSeq stranded mRNA kit following the manufacturer’s instructions (Illumina). Individual libraries were pooled and sequenced on the Illumina HiSeq4000 using the paired-end 150nt mode. Between ∼10-40 million paired end reads were generated per sample and three replicates were sequenced for each timepoint in the desiccation and rehydration experiments for roots and leaves.

### DNA extraction, PacBio, and HiC sequencing

Young leaf tissue of growth chamber grown Craterostigma plants was used for high molecular weight (HMW) genomic DNA and for HiC chromosome conformation capture library preparation. Genomic DNA was isolated using a modified DNA preparation (51) followed by phenol chloroform purification. PacBio libraries were constructed and size selected for ∼25kb fragments on the BluePippen system (Sage Science) and libraries were purified using AMPure XP beads (Beckman Coulter). Libraries were sequenced on a Sequel platform with V4 software and V2 chemistry. In total, 3,590,703 filtered subreads spanning 74.7 Gb were sequenced, and this data corresponds to ∼72 X coverage of the Craterostgma genome. A HiC chromosome conformation capture library was constructed using 0.2 g of fresh, young Craterostigma leaf tissue with the Proximo™ HiC Plant kit (Phase Genomics) following the manufacture’s protocol. The final library was size selected for 300-600 bp and sequenced on the Illumina HiSeq4000 under paired-end 150 bp mode. This HiC data was also used to polish the PacBio-based contigs.

### Assembling the complex Craterostigma genome

Craterostigma plantagineum is octoploid with an estimated haploid genome size of 1.03 Gb based on flow cytometry (55). This corresponds to a monoploid genome size of ∼250 Mb. We assembled the Craterostigma using high coverage (∼74x) PacBio data and scaffolded the resulting contigs using HiC data. The raw PacBio reads were error corrected and assembled using Canu (V2.2) (20). The polished PacBio reads were also assembled with hifiasm (v. 0.15.5) (56) and wtdbg2 (v2.5) (57), but these algorithms generally produced assemblies with lower contiguity or less haplotype-specific sequences. The following Canu parameters were modified, and all others were left as default: minReadLength=5000, GenomeSize=1030mb, minOverlapLength=1000. Pilon (v 1.22) (58) was used with high coverage Illumina data to polish the Canu based contigs and remove residual errors. Illumina reads were quality trimmed using fastp (v 0.23) (59) and aligned to the draft contigs using bowtie2 (v2.3.0)(60) with default parameters. The resulting sorted bam file was used as input for Pilon and the following parameters were modified: --flank 7, --K 49, and --mindepth 15. This process was repeated three times, with Pilon run recursively after each round using the updated reference each time.

Contigs were scaffolded using a Hi-C proximity-based assembly approach as previously described (61). 150 bp paired end Illumina reads were generated for the HiC library to reduce chimeric read mapping across haplotypes and homeologous chromosome regions. The Juicer and 3d-DNA pipelines were used to process and filter the HiC data and for scaffolding (62). HiC reads were quality trimmed using fastp (v 0.23) (59) and aligned to the pacbio-based contigs using BWA (v0.7.16) (63) with strict parameters (-n 0) to prevent chimeric read mapping. Contigs were ordered and scaffolded using the 3d-DNA pipeline (v 180922) with the misassembly identification disabled (-r 0) to avoid erroneous contig splitting due to chimaeric read mapping (64). The resulting HiC contact matrix was visualized using Juicebox, and misjoins and ordering issues were manually corrected based on neighboring interactions. The complex polyploidy and assembly of multiple haplotypes prevented us from generating a chromosome scale assembly, but dozens of full-length chromosomes were scaffolded and validated based on synteny with *L. brevidens*. This scaffolded version (V4) was used for all downstream analyses.

### Genome annotation

The Craterostigma genome was annotated using the MAKER-P pipeline (v2.31.10) (23) with the following sets of input data. Transcript evidence was generated using the desiccation and rehydration RNAseq datasets from root and leaf tissue. RNAseq reads were quality trimmed using fastp (v 0.23) (59) and aligned to the unmasked Craterostigma genome using the splice aware alignment program STAR (v2.6) (65). A set of non-overlapping transcripts were identified using StringTie (v1.3.4) (66) with default parameters, and the resulting gff file was used as transcript evidence for MAKER. A set of protein sequences from *L. brevidens* and *L. subracemosa* (51) as well as Arabidopsis were used as protein evidence. A library of repetitive elements was constructed for Craterostigma using the EDTA package (v2.0.0) (67) which identifies DNA based transposable elements using HelitronScanner (68) and LTR retrotransposons using LTR_FINDER (69) and LTR_HARVEST (70).

These files were used as input for MAKER-P, and ab initio gene prediction was conducted using SNAP(71) and Augustus (v3.0.2) (72) with two rounds of iterative training. Raw gene models from MAKER were filtered to remove any residual repetitive element derived proteins using BLAST with a non-redundant transposase library. The plant-specific (embryophyta_odb9) set of Benchmarking Universal Single-Copy Orthologs, (BUSCO v.2) (73) was used to assess assembly completeness. After filtering, 222,749 high confidence gene models were identified in Craterostigma.

### Comparative genomics analyses

Comparative genomics analyses between Craterostigma and the two Lindernia species *(L. brevidens* and *L. subracemosa*) were conducted using the MCScan toolkit (v1.1) (74) implemented in python [https://github.com/tanghaibao/jcvi/wiki/MCscan-(Python-version)]. Syntenic orthologs were identified across the three species using *L. brevidens* as an anchor given its high-quality chromosome scale assembly. Gene models were aligned using LAST and syntenic blocks were identified with a minimum number of five overlapping syntenic genes. Macrosynteny and microsynteny plots and syntenic block depths were plotted using the python version of MCScan. A file of syntenic orthologs was generated with a ratio of 1:1:8 for *L. brevidens, L. subracemosa*, and Craterostigma respectively, to account for the octoploidy of Craterostigma. This list of syntenic orthologs was used for downstream comparative genomic and gene expression analyses.

### Gene expression analyses

RNAseq data was analyzed using a pipeline developed in the VanBuren lab to trim, quantify, and identify differentially expressed genes (https://github.com/pardojer23/RNAseqV2). Briefly, raw paired end Illumina reads were quality trimmed using fastp (v0.23) (59) with default parameters. Quality filtered reads were pseudo-aligned to the Craterostigma transcripts (gene models) using Salmon (v1.6.0) (75) with the quasi mapping mode. Transcript level estimates were converted to gene level transcript per million counts using the R package tximport (76) and differential gene expression analyses was performed using DEseq2 (77) with the model yij ∼ μ + timepoint + eij. Co-expression networks were constructed and compared for roots and leaves using Clust (v1.12.0) (47). Clust performs optimized consensus clustering to remove spurious associations and can handle multiple heterogeneous datasets with different numbers of samples. Clust was run with the following parameters: ‘-n 101 3 4, -t 0.5’.

Expression levels for syntenic orthologs were used for cross-species comparisons. For Craterostigma, expression levels for all homeologs were summed into a single value for direct comparisons against the single corresponding ortholog in *L. brevidens* and *L. subracemosa*. Genes with no orthology were filtered out from this analysis. Transformed expression values were used as input into Clust (v1.12.0) (47) and the tightness parameter (-t) was changed to 0.2. Principal component analysis and other statistical analyses were implemented in Python using the package scikit learn (78).

## Notes

### Competing Interest Statement

The authors have declared no competing interest.

