## Supplemental Figures/Tables for "The complex octoploid Craterostigma genome and tissue-specific mechanisms underlying desiccation tolerance"

**Supplemental materials**

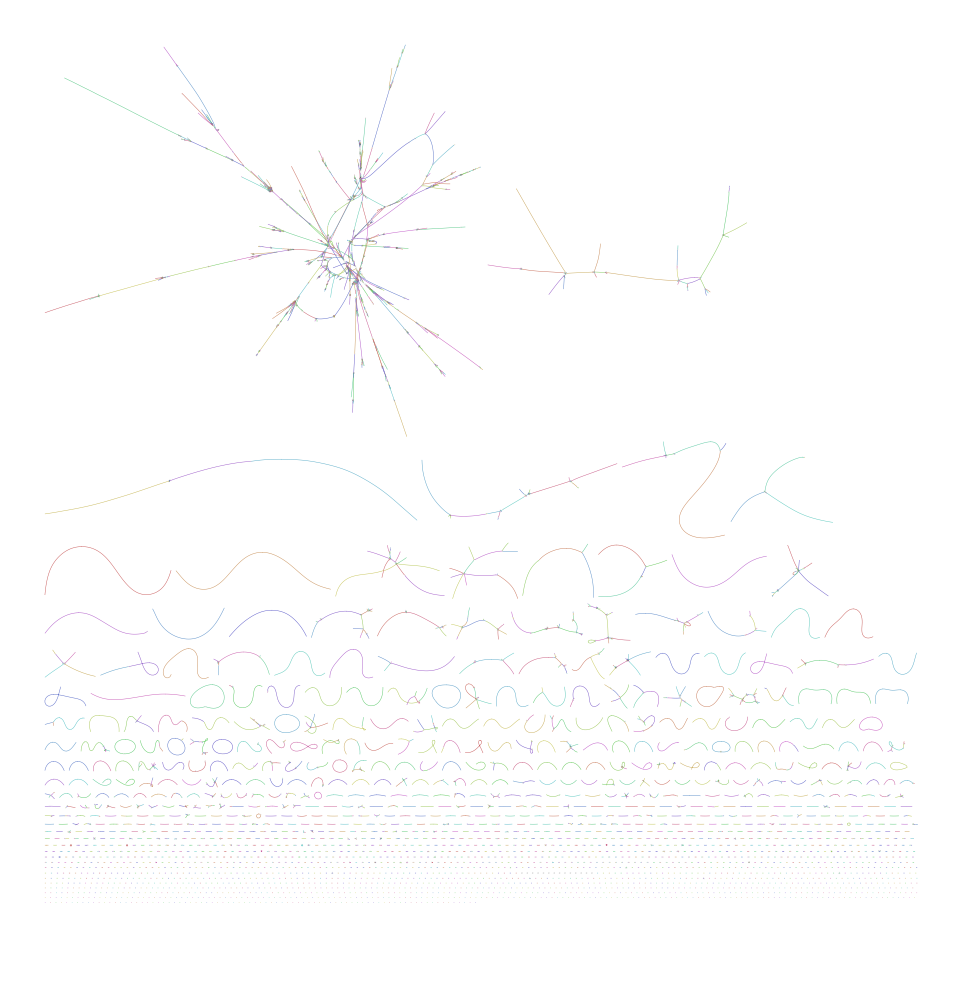

**Supplemental Figure 1. Assembly graph of the Craterostigma genome.** The assembly graph for the V1 genome is plotted where randomly colored nodes (lines) represent individual, contigs, and ambiguities between contigs are shown as connections. Circles within the assembly graph correspond to microbial endophyte genomes that were filtered out prior to HiC scaffolding (See Supplemental Table 2).
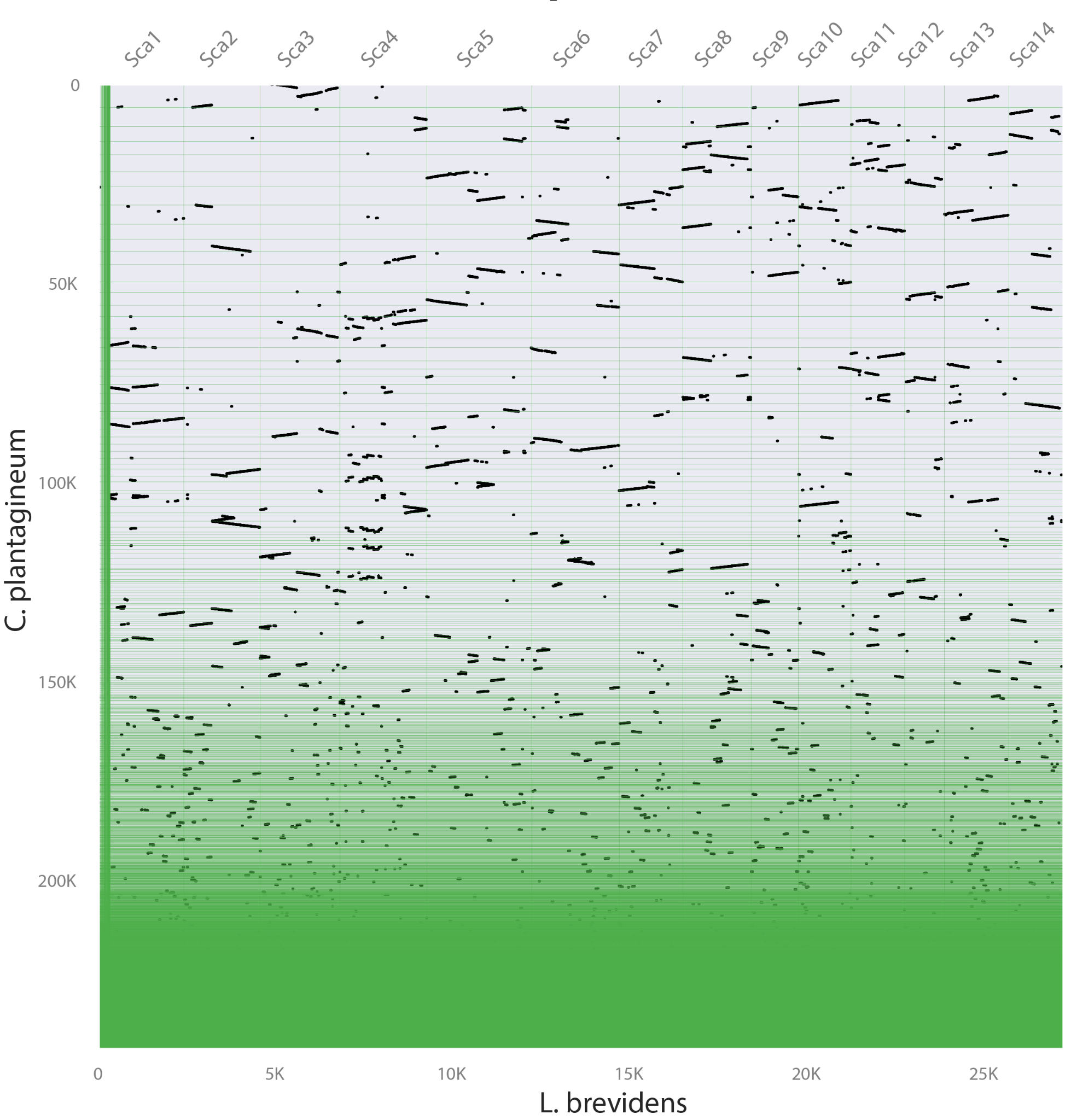

**Supplemental Figure 2. Macrosynteny between the Craterostigma V4 assembly and the *Lindernia brevidens* genome.** The 14 pseudomolecules are plotted for the L. brevidens genome and the draft scaffolds for the Craterostigma V4 genome. Syntenic gene pairs are denoted by black dots, and between 2-8 syntenic regions in Craterostigma are found for each region in *L. brevidens*.

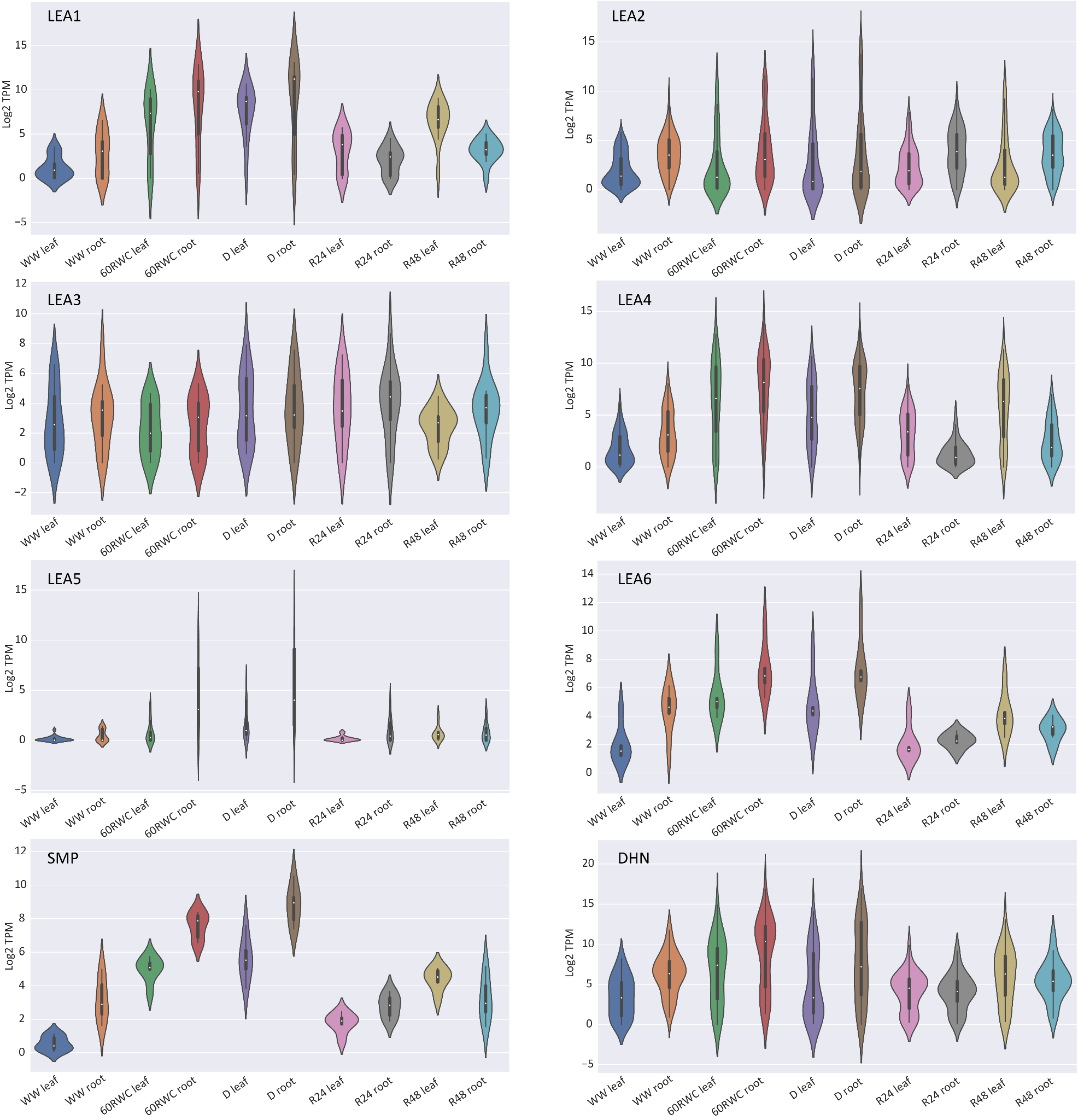

**Supplemental Figure 3. Expression dynamics of LEA family proteins during desiccation and rehydration.** Log2 normalized expression is plotted for various LEA proteins in roots (left) and leaves (right) for the timepoints in the desiccation and rehydration timecourses.

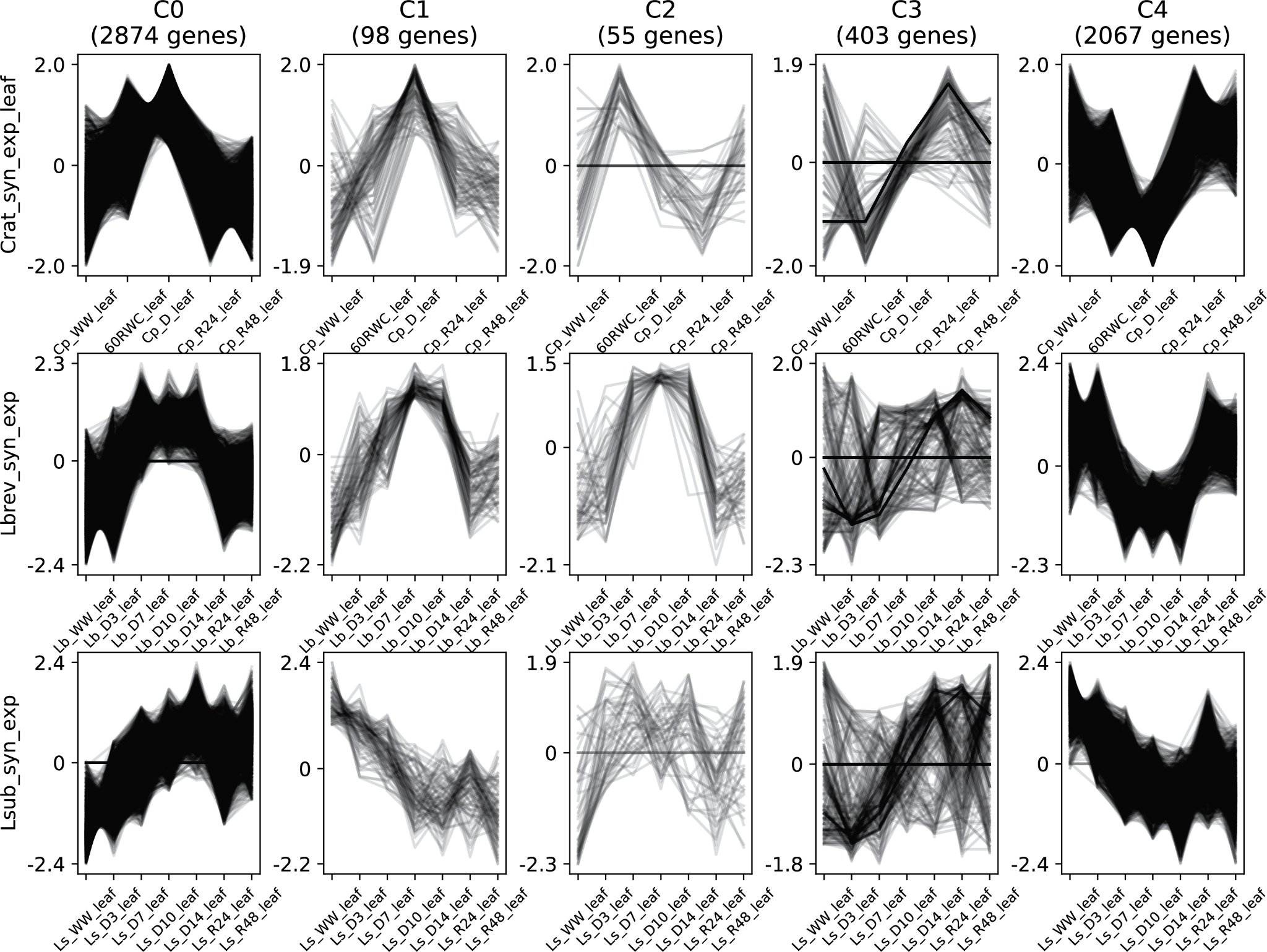

**Supplemental Figure 4. Co-expression of syntenic orthologs across Linderneacaea during desiccation and rehydration timecourses.** Desiccation and rehydration timecourses were collected under the same conditions for all three species and similar (but not identical) timepoints were sampled across species. Genes were clustered into discrete modules using clust with syntenic orthologs between the three species and the transformed expression patterns are plotted for each of the five modules.

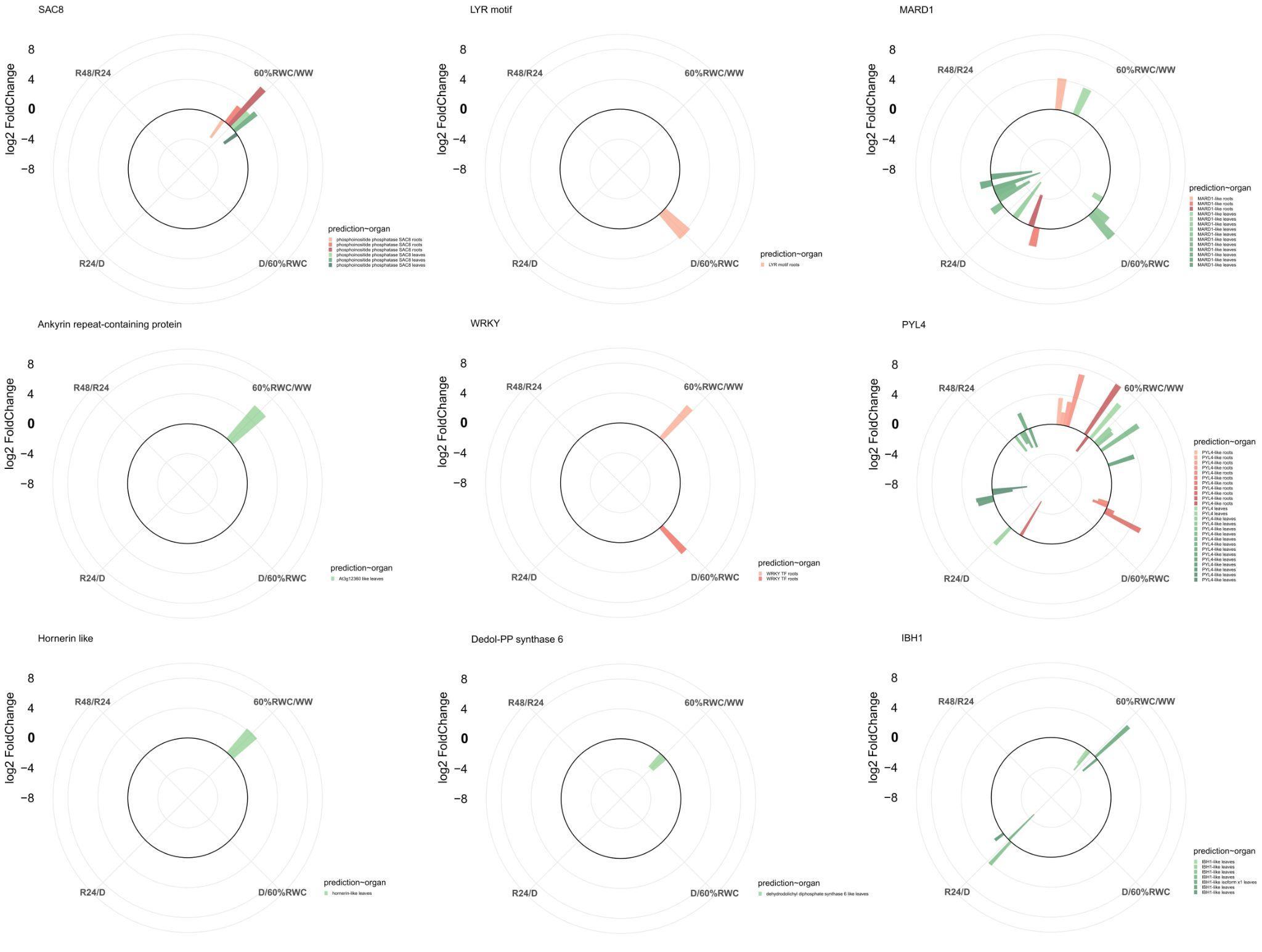

**Supplemental Figure 5. Expression of early responsive genes annotated in significant GO-MF terms during dehydration and desiccation in roots and leaves.** Log2FoldChange expression levels are shown as a contrast between different desiccation and rehydration time courses. WW: well-watered; 60%RWC: moderate desiccation (~60% relative water content); D: dehydration; R24: 24 hours post rehydration; R48: 48 hours post rehydration.

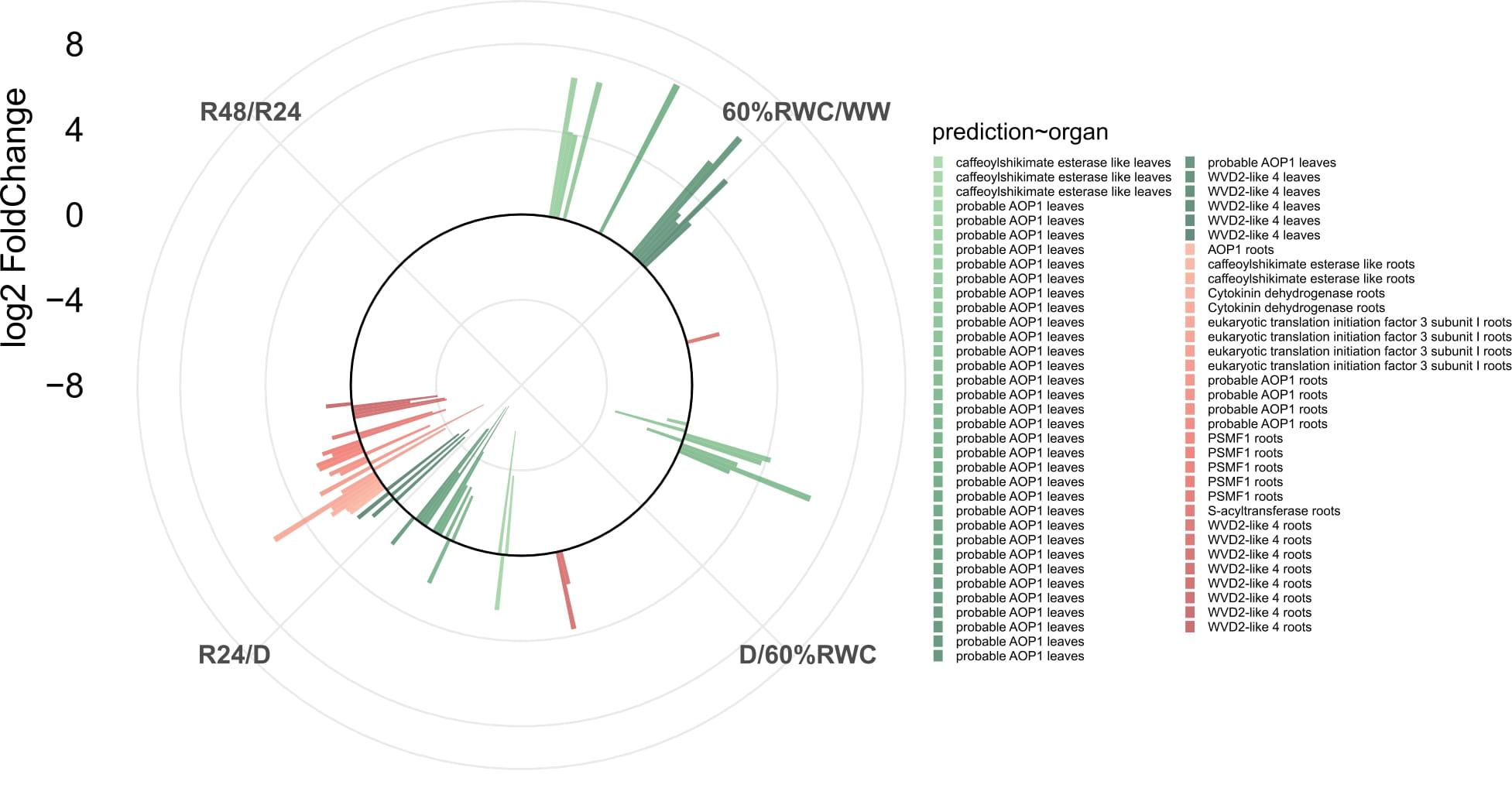

**Supplemental Figure 6. Expression of root associated genes annotated in significant GO-MF terms upon rehydration, compared with leaves expression patterns of the same gene families.** Log2FoldChange expression levels are shown as a contrast between different desiccation and rehydration time courses. WW: well-watered; 60%RWC: moderate desiccation (~60% relative water content); D: dehydration; R24: 24 hours post rehydration; R48: 48 hours post rehydration.

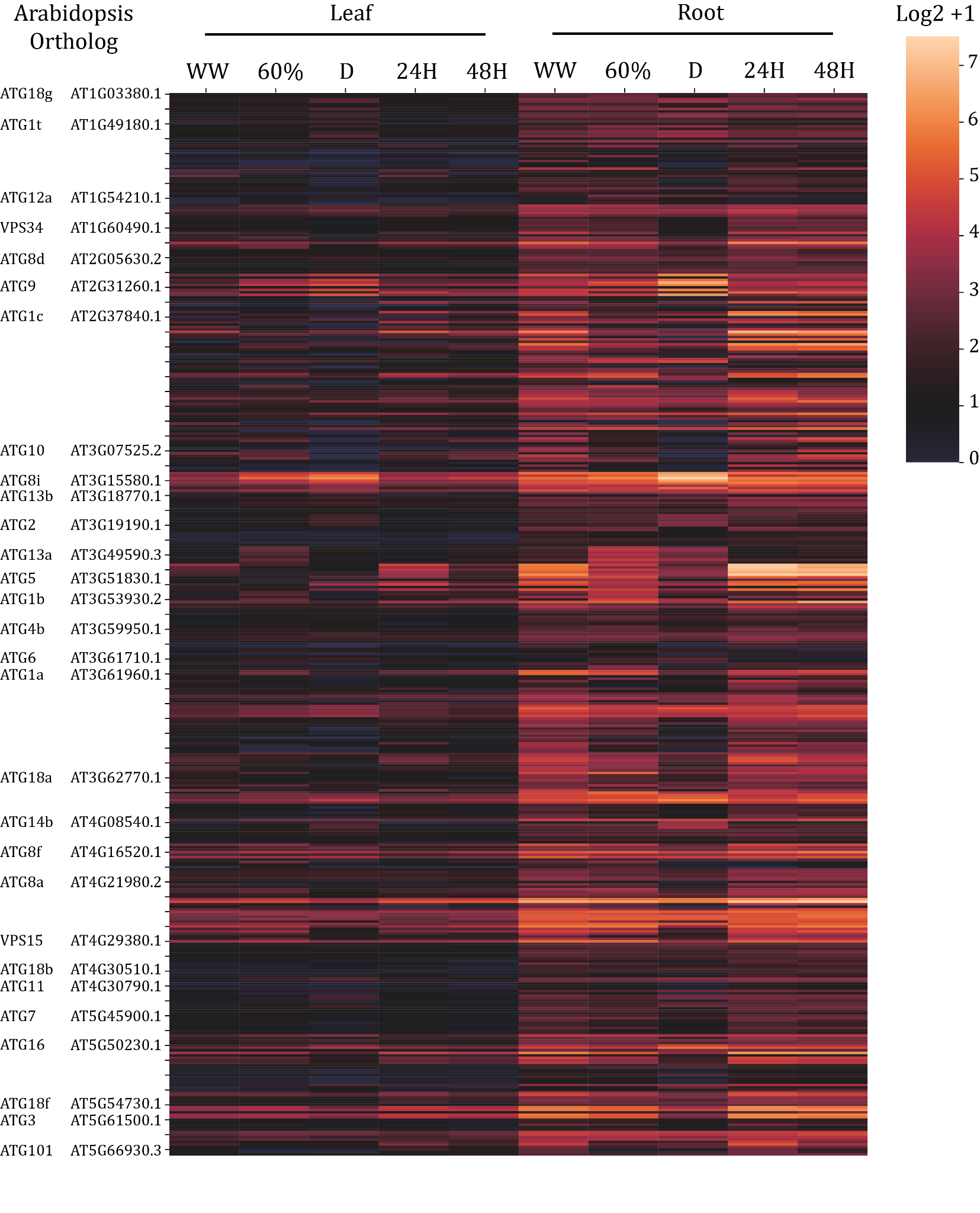

**Supplemental Figure 7. Expression pattern of autophagy associated genes.** A heatmap of Log2 transformed expression data for autophagy associated genes is shown for the leaf and root timecourses. The Arabidopsis and annotated gene names are shown on the left, and the Craterostigma orthologs are plotted in the heatmap.

**Supplemental Table 1. Assembly statistics of the Craterostigma genome.**

|  | **V1** | **V4** |
| --- | --- | --- |
| Contig/scaffold N50 | 1,368,252 | 5,403,000 |
| Numer of contigs/scaffolds | 6,064 | 7,156 |
| Total assembly size | 1,788,238,037 | 1,716,745,304 |

**Supplemental Table 2. Bacterial endophyte genomes in the Craterostigma assembly.**

| **Contig** | **ENA accession** | **Top BLAST hit** | **E value** | **% identity match** |
| --- | --- | --- | --- | --- |
| tig00003525 | CP034121.1 | Microbacterium sp. RG1 | 0 | 85.41% |
| tig00002173 | CP049919.1 | Kinneretia sp. DAIF2 | 0 | 93.65% |
| tig00004026 | CP088918.1 | Rhodanobacter denitrificans | 0 | 89.10% |
| tig00003599 | CP088925.1 | Rhodanobacter denitrificans | 0 | 87.98% |
| tig00009639 | AP010946.1 | Azospirillum sp. B510 | ###### | 75.70% |
| tig00002692 | CP047310.1 | Stenotrophomonas maltophilia | 0 | 98.18% |
| tig00002382 | AM902716.1 | Bordetella petrii | 0 | 83.09% |
| tig00008401 | CP049199.1 | Caulobacter soli | 0 | 79.46% |
| tig00008483 | CP061847.1 | Lysobacter sp. CW239 | ###### | 73.77% |
| tig00000928 | CP074676.1 | Pseudomonas qingdaonensis | 0 | 84.57% |
| tig00009734 | CP045399.1 | Halomonas sp. THAF12 | ###### | 79.38% |
| tig00001362 | LR134361.1 | Achromobacter insolitus | 0 | 82.50% |
| tig00001830 | CP040930.1 | Pseudomonas sp. SWI7 | 0 | 99.76% |
| tig00009852 | CP040750.1 | Plantibacter sp. M259 | ###### | 74.02% |
| tig00003091 | CP084910.1 | Asticcacaulis sp. AND118 | 0 | 80.80% |
| tig00009231 | KP718623.1 | Paulownia coreana | 0 | 94.89% |
| tig00002325 | CP083809.1 | Pantoea agglomerans | 0 | 94.71% |
| tig00003103 | CP010945.1 | Pseudomonas fluorescens NCIMB 11764 | 0 | 97.73% |
| tig00009742 | CP041348.1 | Komagataeibacter xylinus | ###### | 73.81% |
| tig00382919 | CP003555.1 | Advenella kashmirensis WT001 | 0 | 90.05% |
| tig00001504 | CP086100.1 | Enterobacter ludwigii | 0 | 99.19% |
| tig00008970 | CP027260.1 | Pectobacterium parmentieri | 0 | 81.26% |
| tig00002137 | CP040095.1 | Pantoea sp. SO10 | 0 | 89.47% |
| tig00004729 | CP015775.1 | Brucella pseudogrignonensis | 0 | 82.85% |
| tig00005647 | CP022603.1 | [Ochrobactrum] quorumnocens | 0 | 90.94% |
| tig00006626 | CP040096.1 | Pantoea sp. SO10 | 0 | 89.33% |
| tig00007364 | CP044222.1 | Nitrincola iocasae | ###### | 70.87% |
| tig00003666 | CP033953.1 | Methylomonas clara | 0 | 83.24% |
| tig00009638 | CP045103.1 | Acinetobacter johnsonii | 0 | 95.39% |
| tig00001552 | CP031763.1 | Flavobacterium johnsoniae | 0 | 95.78% |
| tig00000787 | CP042170.1 | Flavobacterium sp. KBS0721 | 0 | 82.55% |

**Supplemental Table 3. Shared enriched GO terms upregulated in leaves and roots under desiccation.**

| **GO** | **GO term description** | **p_uncorrected** | **p_fdr_bh** |
| --- | --- | --- | --- |
| GO:0000209 | protein polyubiquitination | 6.44E-08 | 6.04E-05 |
| GO:0000304 | response to singlet oxygen | 1.16E-07 | 6.04E-05 |
| GO:0009450 | gamma-aminobutyric acid catabolic process | 3.63E-07 | 6.04E-05 |
| GO:0009448 | gamma-aminobutyric acid metabolic process | 3.63E-07 | 6.04E-05 |
| GO:0006526 | arginine biosynthetic process | 3.84E-07 | 6.04E-05 |
| GO:0051052 | regulation of DNA metabolic process | 4.46E-07 | 6.04E-05 |
| GO:0010286 | heat acclimation | 5.11E-07 | 6.04E-05 |
| GO:1903409 | reactive oxygen species biosynthetic process | 5.13E-07 | 6.04E-05 |
| GO:0040014 | regulation of multicellular organism growth | 5.22E-07 | 6.04E-05 |
| GO:0010230 | alternative respiration | 5.22E-07 | 6.04E-05 |
| GO:0006572 | tyrosine catabolic process | 5.22E-07 | 6.04E-05 |
| GO:0009649 | entrainment of circadian clock | 5.34E-07 | 6.04E-05 |
| GO:0010099 | regulation of photomorphogenesis | 5.39E-07 | 6.04E-05 |
| GO:2000030 | regulation of response to red or far red light | 5.39E-07 | 6.04E-05 |
| GO:0010100 | negative regulation of photomorphogenesis | 5.93E-07 | 6.04E-05 |
| GO:0000956 | nuclear-transcribed mRNA catabolic process | 5.93E-07 | 6.04E-05 |
| GO:0000018 | regulation of DNA recombination | 6.28E-07 | 6.04E-05 |
| GO:0046786 | viral replication complex formation and maintenance | 6.28E-07 | 6.04E-05 |
| GO:0051053 | negative regulation of DNA metabolic process | 6.28E-07 | 6.04E-05 |
| GO:0045910 | negative regulation of DNA recombination | 6.28E-07 | 6.04E-05 |
| GO:0010498 | proteasomal protein catabolic process | 6.49E-07 | 6.04E-05 |
| GO:0009082 | branched-chain amino acid biosynthetic process | 6.9E-07 | 6.04E-05 |
| GO:0016559 | peroxisome fission | 6.9E-07 | 6.04E-05 |
| GO:0006525 | arginine metabolic process | 6.91E-07 | 6.04E-05 |
| GO:0072663 | establishment of protein localization to peroxisome | 7.06E-07 | 6.04E-05 |
| GO:0006625 | protein targeting to peroxisome | 7.06E-07 | 6.04E-05 |
| GO:0043574 | peroxisomal transport | 7.06E-07 | 6.04E-05 |
| GO:0072662 | protein localization to peroxisome | 7.06E-07 | 6.04E-05 |
| GO:0006402 | mRNA catabolic process | 7.06E-07 | 6.04E-05 |
| GO:0042752 | regulation of circadian rhythm | 7.32E-07 | 6.04E-05 |
| GO:0001666 | response to hypoxia | 7.64E-07 | 6.04E-05 |
| GO:0036293 | response to decreased oxygen levels | 7.64E-07 | 6.04E-05 |
| GO:0070482 | response to oxygen levels | 7.64E-07 | 6.04E-05 |
| GO:0000184 | nuclear-transcribed mRNA catabolic process, nonsense-mediated decay | 7.77E-07 | 6.04E-05 |
| GO:0006419 | alanyl-tRNA aminoacylation | 7.87E-07 | 6.04E-05 |
| GO:0006527 | arginine catabolic process | 7.87E-07 | 6.04E-05 |
| GO:0006540 | glutamate decarboxylation to succinate | 7.87E-07 | 6.04E-05 |
| GO:0006471 | protein ADP-ribosylation | 8.16E-07 | 6.04E-05 |
| GO:0043153 | entrainment of circadian clock by photoperiod | 8.35E-07 | 6.04E-05 |
| GO:0043631 | RNA polyadenylation | 8.88E-07 | 6.04E-05 |
| GO:0006366 | transcription by RNA polymerase II | 8.94E-07 | 6.04E-05 |
| GO:0006401 | RNA catabolic process | 9.31E-07 | 6.04E-05 |
| GO:0009269 | response to desiccation | 9.5E-07 | 6.04E-05 |
| GO:0032984 | protein-containing complex disassembly | 1.05E-06 | 6.04E-05 |
| GO:0043624 | cellular protein complex disassembly | 1.1E-06 | 6.04E-05 |
| GO:0006415 | translational termination | 1.1E-06 | 6.04E-05 |
| GO:0007140 | male meiotic nuclear division | 1.12E-06 | 6.04E-05 |
| GO:0140013 | meiotic nuclear division | 1.12E-06 | 6.04E-05 |
| GO:0000305 | response to oxygen radical | 1.17E-06 | 6.04E-05 |
| GO:0000303 | response to superoxide | 1.17E-06 | 6.04E-05 |
| GO:0010182 | sugar mediated signaling pathway | 1.17E-06 | 6.04E-05 |
| GO:0009756 | carbohydrate mediated signaling | 1.17E-06 | 6.04E-05 |
| GO:0006566 | threonine metabolic process | 1.21E-06 | 6.04E-05 |
| GO:0043248 | proteasome assembly | 1.23E-06 | 6.04E-05 |
| GO:0051788 | response to misfolded protein | 1.23E-06 | 6.04E-05 |
| GO:0031087 | deadenylation-independent decapping of nuclear-transcribed mRNA | 1.26E-06 | 6.04E-05 |
| GO:0110154 | RNA decapping | 1.26E-06 | 6.04E-05 |
| GO:0110156 | methylguanosine-cap decapping | 1.26E-06 | 6.04E-05 |
| GO:0009081 | branched-chain amino acid metabolic process | 1.29E-06 | 6.04E-05 |
| GO:0006744 | ubiquinone biosynthetic process | 1.32E-06 | 6.04E-05 |
| GO:0006743 | ubiquinone metabolic process | 1.32E-06 | 6.04E-05 |
| GO:0006289 | nucleotide-excision repair | 1.32E-06 | 6.04E-05 |
| GO:0009062 | fatty acid catabolic process | 1.35E-06 | 6.04E-05 |
| GO:0006635 | fatty acid beta-oxidation | 1.39E-06 | 6.04E-05 |
| GO:0007031 | peroxisome organization | 1.41E-06 | 6.04E-05 |
| GO:0072593 | reactive oxygen species metabolic process | 1.41E-06 | 6.04E-05 |
| GO:0016226 | iron-sulfur cluster assembly | 1.43E-06 | 6.04E-05 |
| GO:0031163 | metallo-sulfur cluster assembly | 1.43E-06 | 6.04E-05 |
| GO:0015980 | energy derivation by oxidation of organic compounds | 1.48E-06 | 6.04E-05 |
| GO:0030258 | lipid modification | 1.49E-06 | 6.04E-05 |
| GO:0072598 | protein localization to chloroplast | 1.56E-06 | 6.04E-05 |
| GO:0045036 | protein targeting to chloroplast | 1.56E-06 | 6.04E-05 |
| GO:0072596 | establishment of protein localization to chloroplast | 1.56E-06 | 6.04E-05 |
| GO:0043161 | proteasome-mediated ubiquitin-dependent protein catabolic process | 1.56E-06 | 6.04E-05 |
| GO:0050792 | regulation of viral process | 1.64E-06 | 6.04E-05 |
| GO:0015985 | energy coupled proton transport, down electrochemical gradient | 1.65E-06 | 6.04E-05 |
| GO:0015986 | ATP synthesis coupled proton transport | 1.65E-06 | 6.04E-05 |
| GO:0000377 | RNA splicing, via transesterification reactions with bulged adenosine as nucleophile | 1.73E-06 | 6.04E-05 |
| GO:0009064 | glutamine family amino acid metabolic process | 1.73E-06 | 6.04E-05 |
| GO:0000375 | RNA splicing, via transesterification reactions | 1.73E-06 | 6.04E-05 |
| GO:0009112 | nucleobase metabolic process | 1.79E-06 | 6.08E-05 |
| GO:0019395 | fatty acid oxidation | 1.8E-06 | 6.08E-05 |
| GO:0034440 | lipid oxidation | 1.8E-06 | 6.08E-05 |
| GO:0035194 | post-transcriptional gene silencing by RNA | 1.9E-06 | 6.24E-05 |
| GO:0065002 | intracellular protein transmembrane transport | 1.9E-06 | 6.24E-05 |
| GO:0071806 | protein transmembrane transport | 1.9E-06 | 6.24E-05 |
| GO:0009065 | glutamine family amino acid catabolic process | 1.93E-06 | 6.27E-05 |
| GO:0007005 | mitochondrion organization | 2.01E-06 | 6.3E-05 |
| GO:0008380 | RNA splicing | 2.01E-06 | 6.3E-05 |
| GO:1901606 | alpha-amino acid catabolic process | 2.02E-06 | 6.3E-05 |
| GO:0030163 | protein catabolic process | 2.09E-06 | 6.36E-05 |
| GO:0000398 | mRNA splicing, via spliceosome | 2.1E-06 | 6.36E-05 |
| GO:0009063 | cellular amino acid catabolic process | 2.13E-06 | 6.4E-05 |
| GO:0006352 | DNA-templated transcription, initiation | 2.18E-06 | 6.43E-05 |
| GO:0072329 | monocarboxylic acid catabolic process | 2.21E-06 | 6.43E-05 |
| GO:0009644 | response to high light intensity | 2.21E-06 | 6.43E-05 |
| GO:0034470 | ncRNA processing | 2.48E-06 | 6.43E-05 |
| GO:0072594 | establishment of protein localization to organelle | 2.48E-06 | 6.43E-05 |
| GO:0007059 | chromosome segregation | 2.52E-06 | 6.43E-05 |
| GO:0046395 | carboxylic acid catabolic process | 2.53E-06 | 6.43E-05 |
| GO:0048285 | organelle fission | 2.55E-06 | 6.43E-05 |
| GO:0006839 | mitochondrial transport | 2.56E-06 | 6.43E-05 |
| GO:0006397 | mRNA processing | 2.61E-06 | 6.43E-05 |
| GO:0072655 | establishment of protein localization to mitochondrion | 2.64E-06 | 6.43E-05 |
| GO:0070585 | protein localization to mitochondrion | 2.64E-06 | 6.43E-05 |
| GO:0006413 | translational initiation | 2.67E-06 | 6.43E-05 |
| GO:0016071 | mRNA metabolic process | 2.67E-06 | 6.43E-05 |
| GO:0009642 | response to light intensity | 2.92E-06 | 6.85E-05 |
| GO:0009657 | plastid organization | 2.98E-06 | 6.93E-05 |
| GO:0097659 | nucleic acid-templated transcription | 3.01E-06 | 6.93E-05 |
| GO:0006351 | transcription, DNA-templated | 3.01E-06 | 6.93E-05 |
| GO:0070647 | protein modification by small protein conjugation or removal | 3.14E-06 | 7.09E-05 |
| GO:0035966 | response to topologically incorrect protein | 3.22E-06 | 7.09E-05 |
| GO:0044282 | small molecule catabolic process | 3.3E-06 | 7.09E-05 |
| GO:0033365 | protein localization to organelle | 3.32E-06 | 7.09E-05 |
| GO:1901565 | organonitrogen compound catabolic process | 3.33E-06 | 7.09E-05 |
| GO:0006281 | DNA repair | 3.34E-06 | 7.09E-05 |
| GO:0006593 | ornithine catabolic process | 3.37E-06 | 7.09E-05 |
| GO:0006862 | nucleotide transport | 3.38E-06 | 7.09E-05 |
| GO:0000278 | mitotic cell cycle | 3.38E-06 | 7.09E-05 |
| GO:0016054 | organic acid catabolic process | 3.42E-06 | 7.09E-05 |
| GO:0034613 | cellular protein localization | 3.43E-06 | 7.09E-05 |
| GO:0009658 | chloroplast organization | 3.45E-06 | 7.09E-05 |
| GO:0006470 | protein dephosphorylation | 3.54E-06 | 7.09E-05 |
| GO:0032774 | RNA biosynthetic process | 3.55E-06 | 7.09E-05 |
| GO:0006605 | protein targeting | 3.56E-06 | 7.09E-05 |
| GO:0006567 | threonine catabolic process | 3.57E-06 | 7.09E-05 |
| GO:0070727 | cellular macromolecule localization | 3.72E-06 | 7.13E-05 |
| GO:0045333 | cellular respiration | 3.85E-06 | 7.13E-05 |
| GO:0006888 | endoplasmic reticulum to Golgi vesicle-mediated transport | 4.02E-06 | 7.13E-05 |
| GO:0001101 | response to acid chemical | 4.09E-06 | 7.13E-05 |
| GO:0022411 | cellular component disassembly | 4.16E-06 | 7.13E-05 |
| GO:0016567 | protein ubiquitination | 4.17E-06 | 7.13E-05 |
| GO:0042023 | DNA endoreduplication | 4.2E-06 | 7.13E-05 |
| GO:0009408 | response to heat | 4.21E-06 | 7.13E-05 |
| GO:0031123 | RNA 3'-end processing | 4.4E-06 | 7.13E-05 |
| GO:0010155 | regulation of proton transport | 4.7E-06 | 7.27E-05 |
| GO:1904062 | regulation of cation transmembrane transport | 4.7E-06 | 7.27E-05 |
| GO:0016192 | vesicle-mediated transport | 4.76E-06 | 7.33E-05 |
| GO:0044265 | cellular macromolecule catabolic process | 4.94E-06 | 7.44E-05 |
| GO:0009251 | glucan catabolic process | 4.98E-06 | 7.44E-05 |
| GO:0044247 | cellular polysaccharide catabolic process | 4.98E-06 | 7.44E-05 |
| GO:0048581 | negative regulation of post-embryonic development | 5E-06 | 7.44E-05 |
| GO:1901605 | alpha-amino acid metabolic process | 5.08E-06 | 7.44E-05 |
| GO:0006511 | ubiquitin-dependent protein catabolic process | 5.13E-06 | 7.44E-05 |
| GO:0019941 | modification-dependent protein catabolic process | 5.13E-06 | 7.44E-05 |
| GO:0043632 | modification-dependent macromolecule catabolic process | 5.13E-06 | 7.44E-05 |
| GO:0006457 | protein folding | 5.25E-06 | 7.52E-05 |
| GO:0009415 | response to water | 5.28E-06 | 7.52E-05 |
| GO:0051603 | proteolysis involved in cellular protein catabolic process | 5.28E-06 | 7.52E-05 |
| GO:0017038 | protein import | 5.29E-06 | 7.52E-05 |
| GO:0006396 | RNA processing | 5.33E-06 | 7.54E-05 |
| GO:0006886 | intracellular protein transport | 5.38E-06 | 7.58E-05 |
| GO:0044242 | cellular lipid catabolic process | 5.52E-06 | 7.75E-05 |
| GO:0048519 | negative regulation of biological process | 5.57E-06 | 7.75E-05 |
| GO:0006996 | organelle organization | 5.63E-06 | 7.81E-05 |
| GO:0034654 | nucleobase-containing compound biosynthetic process | 5.75E-06 | 7.84E-05 |
| GO:0051928 | positive regulation of calcium ion transport | 5.82E-06 | 7.84E-05 |
| GO:0010959 | regulation of metal ion transport | 5.82E-06 | 7.84E-05 |
| GO:0051924 | regulation of calcium ion transport | 5.82E-06 | 7.84E-05 |
| GO:0055114 | oxidation-reduction process | 5.91E-06 | 7.92E-05 |
| GO:0008104 | protein localization | 6.12E-06 | 8.11E-05 |
| GO:0009793 | embryo development ending in seed dormancy | 6.2E-06 | 8.12E-05 |
| GO:0010038 | response to metal ion | 6.24E-06 | 8.13E-05 |
| GO:0009790 | embryo development | 6.26E-06 | 8.13E-05 |
| GO:0009057 | macromolecule catabolic process | 6.33E-06 | 8.18E-05 |
| GO:0015031 | protein transport | 6.38E-06 | 8.22E-05 |
| GO:0051649 | establishment of localization in cell | 6.42E-06 | 8.24E-05 |
| GO:0016444 | somatic cell DNA recombination | 6.51E-06 | 8.32E-05 |
| GO:0048580 | regulation of post-embryonic development | 6.58E-06 | 8.35E-05 |
| GO:0045184 | establishment of protein localization | 6.66E-06 | 8.36E-05 |
| GO:0018130 | heterocycle biosynthetic process | 6.69E-06 | 8.36E-05 |
| GO:0046907 | intracellular transport | 6.72E-06 | 8.37E-05 |
| GO:0033036 | macromolecule localization | 6.78E-06 | 8.38E-05 |
| GO:0006974 | cellular response to DNA damage stimulus | 6.85E-06 | 8.43E-05 |
| GO:2000026 | regulation of multicellular organismal development | 6.95E-06 | 8.51E-05 |
| GO:0051641 | cellular localization | 7.01E-06 | 8.51E-05 |
| GO:0015833 | peptide transport | 7.01E-06 | 8.51E-05 |
| GO:0046686 | response to cadmium ion | 7.1E-06 | 8.58E-05 |
| GO:0016070 | RNA metabolic process | 7.14E-06 | 8.59E-05 |
| GO:0015865 | purine nucleotide transport | 7.2E-06 | 8.64E-05 |
| GO:0009892 | negative regulation of metabolic process | 7.32E-06 | 8.75E-05 |
| GO:0000373 | Group II intron splicing | 7.38E-06 | 8.76E-05 |
| GO:0042886 | amide transport | 7.52E-06 | 8.84E-05 |
| GO:0010016 | shoot system morphogenesis | 7.53E-06 | 8.84E-05 |
| GO:0009266 | response to temperature stimulus | 7.59E-06 | 8.87E-05 |
| GO:0006970 | response to osmotic stress | 7.61E-06 | 8.87E-05 |
| GO:0044248 | cellular catabolic process | 7.64E-06 | 8.88E-05 |
| GO:0009718 | anthocyanin-containing compound biosynthetic process | 7.78E-06 | 8.97E-05 |
| GO:0007275 | multicellular organism development | 8.06E-06 | 9.24E-05 |
| GO:0009651 | response to salt stress | 8.07E-06 | 9.24E-05 |
| GO:0071705 | nitrogen compound transport | 8.2E-06 | 9.36E-05 |
| GO:0051239 | regulation of multicellular organismal process | 8.26E-06 | 9.39E-05 |
| GO:1901575 | organic substance catabolic process | 8.34E-06 | 9.46E-05 |
| GO:0006508 | proteolysis | 8.48E-06 | 9.57E-05 |
| GO:0009084 | glutamine family amino acid biosynthetic process | 8.75E-06 | 9.75E-05 |
| GO:0010087 | phloem or xylem histogenesis | 8.81E-06 | 9.75E-05 |
| GO:0009056 | catabolic process | 8.86E-06 | 9.78E-05 |
| GO:0010035 | response to inorganic substance | 9.03E-06 | 9.89E-05 |
| GO:0090304 | nucleic acid metabolic process | 9.4E-06 | 0.000102 |
| GO:0006520 | cellular amino acid metabolic process | 9.78E-06 | 0.000106 |
| GO:0009414 | response to water deprivation | 9.94E-06 | 0.000107 |
| GO:0006265 | DNA topological change | 1.01E-05 | 0.000108 |
| GO:0003006 | developmental process involved in reproduction | 1.01E-05 | 0.000108 |
| GO:0006139 | nucleobase-containing compound metabolic process | 1.05E-05 | 0.000112 |
| GO:0031047 | gene silencing by RNA | 1.08E-05 | 0.000114 |
| GO:0007530 | sex determination | 1.12E-05 | 0.000116 |
| GO:0035266 | meristem growth | 1.12E-05 | 0.000116 |
| GO:0006435 | threonyl-tRNA aminoacylation | 1.12E-05 | 0.000116 |
| GO:0009409 | response to cold | 1.13E-05 | 0.000117 |
| GO:0022414 | reproductive process | 1.14E-05 | 0.000117 |
| GO:0046483 | heterocycle metabolic process | 1.15E-05 | 0.000118 |
| GO:0006725 | cellular aromatic compound metabolic process | 1.15E-05 | 0.000118 |
| GO:0032501 | multicellular organismal process | 1.2E-05 | 0.000122 |
| GO:0048856 | anatomical structure development | 1.21E-05 | 0.000123 |
| GO:0051054 | positive regulation of DNA metabolic process | 1.25E-05 | 0.000125 |
| GO:0045739 | positive regulation of DNA repair | 1.25E-05 | 0.000125 |
| GO:2001022 | positive regulation of response to DNA damage stimulus | 1.25E-05 | 0.000125 |
| GO:0034641 | cellular nitrogen compound metabolic process | 1.27E-05 | 0.000127 |
| GO:1901360 | organic cyclic compound metabolic process | 1.29E-05 | 0.000128 |
| GO:0009628 | response to abiotic stimulus | 1.31E-05 | 0.000129 |
| GO:0046283 | anthocyanin-containing compound metabolic process | 1.39E-05 | 0.000133 |
| GO:0006624 | vacuolar protein processing | 1.51E-05 | 0.00014 |
| GO:0045694 | regulation of embryo sac egg cell differentiation | 1.51E-05 | 0.00014 |
| GO:0031124 | mRNA 3'-end processing | 1.53E-05 | 0.000141 |
| GO:0006378 | mRNA polyadenylation | 1.53E-05 | 0.000141 |
| GO:0009060 | aerobic respiration | 1.72E-05 | 0.000157 |
| GO:0010555 | response to mannitol | 1.72E-05 | 0.000157 |
| GO:0031146 | SCF-dependent proteasomal ubiquitin-dependent protein catabolic process | 1.78E-05 | 0.000161 |
| GO:0016311 | dephosphorylation | 1.84E-05 | 0.000165 |
| GO:0019752 | carboxylic acid metabolic process | 1.92E-05 | 0.000171 |
| GO:0044786 | cell cycle DNA replication | 1.92E-05 | 0.000171 |
| GO:0016042 | lipid catabolic process | 2E-05 | 0.000177 |
| GO:0006807 | nitrogen compound metabolic process | 2.11E-05 | 0.000185 |
| GO:0044237 | cellular metabolic process | 2.2E-05 | 0.000191 |
| GO:0048439 | flower morphogenesis | 2.26E-05 | 0.000196 |
| GO:0009636 | response to toxic substance | 2.41E-05 | 0.000208 |
| GO:0016043 | cellular component organization | 2.56E-05 | 0.000221 |
| GO:0009987 | cellular process | 2.57E-05 | 0.000221 |
| GO:0051703 | biological process involved in intraspecies interaction between organisms | 2.85E-05 | 0.000245 |
| GO:0006367 | transcription initiation from RNA polymerase II promoter | 2.91E-05 | 0.000247 |
| GO:0016072 | rRNA metabolic process | 2.93E-05 | 0.000247 |
| GO:0006364 | rRNA processing | 2.93E-05 | 0.000247 |
| GO:0019438 | aromatic compound biosynthetic process | 3.14E-05 | 0.000261 |
| GO:0006353 | DNA-templated transcription, termination | 3.23E-05 | 0.000265 |
| GO:0051167 | xylulose 5-phosphate metabolic process | 3.23E-05 | 0.000265 |
| GO:0019649 | formaldehyde assimilation | 3.23E-05 | 0.000265 |
| GO:0015675 | nickel cation transport | 3.23E-05 | 0.000265 |
| GO:0019648 | formaldehyde assimilation via xylulose monophosphate cycle | 3.23E-05 | 0.000265 |
| GO:0048645 | animal organ formation | 3.28E-05 | 0.000269 |
| GO:0043903 | regulation of biological process involved in symbiotic interaction | 3.5E-05 | 0.000285 |
| GO:0046364 | monosaccharide biosynthetic process | 3.75E-05 | 0.000304 |
| GO:0032446 | protein modification by small protein conjugation | 3.91E-05 | 0.000315 |
| GO:0046700 | heterocycle catabolic process | 4.56E-05 | 0.000366 |
| GO:0006261 | DNA-dependent DNA replication | 4.74E-05 | 0.000379 |
| GO:0090630 | activation of GTPase activity | 4.85E-05 | 0.000384 |
| GO:0043547 | positive regulation of GTPase activity | 4.85E-05 | 0.000384 |
| GO:0051345 | positive regulation of hydrolase activity | 4.85E-05 | 0.000384 |
| GO:0045037 | protein import into chloroplast stroma | 5.05E-05 | 0.000398 |
| GO:0009757 | hexose mediated signaling | 5.05E-05 | 0.000398 |
| GO:0007062 | sister chromatid cohesion | 5.1E-05 | 0.000401 |
| GO:0010205 | photoinhibition | 5.38E-05 | 0.000421 |
| GO:1905156 | negative regulation of photosynthesis | 5.38E-05 | 0.000421 |
| GO:0043155 | negative regulation of photosynthesis, light reaction | 5.38E-05 | 0.000421 |
| GO:0010183 | pollen tube guidance | 5.68E-05 | 0.000439 |
| GO:0042330 | taxis | 5.68E-05 | 0.000439 |
| GO:0040011 | locomotion | 5.68E-05 | 0.000439 |
| GO:0050918 | positive chemotaxis | 5.68E-05 | 0.000439 |
| GO:0006935 | chemotaxis | 5.68E-05 | 0.000439 |
| GO:0051241 | negative regulation of multicellular organismal process | 5.8E-05 | 0.000447 |
| GO:0034660 | ncRNA metabolic process | 5.82E-05 | 0.000447 |
| GO:0010265 | SCF complex assembly | 5.9E-05 | 0.000452 |
| GO:0009910 | negative regulation of flower development | 5.93E-05 | 0.000453 |
| GO:0006801 | superoxide metabolic process | 6.62E-05 | 0.000498 |
| GO:0051168 | nuclear export | 6.62E-05 | 0.000498 |
| GO:0006144 | purine nucleobase metabolic process | 6.62E-05 | 0.000498 |
| GO:0010605 | negative regulation of macromolecule metabolic process | 7.15E-05 | 0.000536 |
| GO:0006809 | nitric oxide biosynthetic process | 7.92E-05 | 0.000588 |
| GO:0046209 | nitric oxide metabolic process | 7.92E-05 | 0.000588 |
| GO:0010111 | glyoxysome organization | 7.97E-05 | 0.00059 |
| GO:0016032 | viral process | 8.08E-05 | 0.000597 |
| GO:0006767 | water-soluble vitamin metabolic process | 8.86E-05 | 0.000649 |
| GO:0009631 | cold acclimation | 0.000103 | 0.000737 |
| GO:0005983 | starch catabolic process | 0.000103 | 0.000737 |
| GO:0019660 | glycolytic fermentation | 0.000105 | 0.000737 |
| GO:0042554 | superoxide anion generation | 0.000105 | 0.000737 |
| GO:0046113 | nucleobase catabolic process | 0.000105 | 0.000737 |
| GO:0019650 | glycolytic fermentation to butanediol | 0.000105 | 0.000737 |
| GO:0006554 | lysine catabolic process | 0.000105 | 0.000737 |
| GO:0022616 | DNA strand elongation | 0.000105 | 0.000737 |
| GO:0006145 | purine nucleobase catabolic process | 0.000105 | 0.000737 |
| GO:0006271 | DNA strand elongation involved in DNA replication | 0.000105 | 0.000737 |
| GO:0046110 | xanthine metabolic process | 0.000105 | 0.000737 |
| GO:0019477 | L-lysine catabolic process | 0.000105 | 0.000737 |
| GO:0072523 | purine-containing compound catabolic process | 0.000105 | 0.000737 |
| GO:0046440 | L-lysine metabolic process | 0.000105 | 0.000737 |
| GO:0010089 | xylem development | 0.000107 | 0.000755 |
| GO:0009648 | photoperiodism | 0.000111 | 0.000781 |
| GO:0006885 | regulation of pH | 0.000113 | 0.00079 |
| GO:0010104 | regulation of ethylene-activated signaling pathway | 0.000121 | 0.000838 |
| GO:0070297 | regulation of phosphorelay signal transduction system | 0.000121 | 0.000838 |
| GO:0071702 | organic substance transport | 0.000121 | 0.000838 |
| GO:0034220 | ion transmembrane transport | 0.000136 | 0.000934 |
| GO:0015868 | purine ribonucleotide transport | 0.000138 | 0.000948 |
| GO:0000266 | mitochondrial fission | 0.000138 | 0.000949 |
| GO:2000242 | negative regulation of reproductive process | 0.000139 | 0.00095 |
| GO:0046473 | phosphatidic acid metabolic process | 0.000141 | 0.000957 |
| GO:0043436 | oxoacid metabolic process | 0.000142 | 0.000962 |
| GO:0015931 | nucleobase-containing compound transport | 0.000144 | 0.000974 |

**Supplemental Table 4. Enriched GO terms upregulated in only leaves under desiccation.**

| **GO** | **GO term description** | **p_uncorrected** | **p_fdr_bh** |
| --- | --- | --- | --- |
| GO:0090501 | RNA phosphodiester bond hydrolysis | 2.38E-07 | 0.000126 |
| GO:0044743 | protein transmembrane import into intracellular organelle | 2.79E-07 | 0.000126 |
| GO:0090305 | nucleic acid phosphodiester bond hydrolysis | 3E-07 | 0.000126 |
| GO:0010497 | plasmodesmata-mediated intercellular transport | 3.58E-07 | 0.000126 |
| GO:0010496 | intercellular transport | 3.58E-07 | 0.000126 |
| GO:0042793 | plastid transcription | 4.07E-07 | 0.000126 |
| GO:0006364 | rRNA processing | 4.55E-07 | 0.000126 |
| GO:0016072 | rRNA metabolic process | 4.55E-07 | 0.000126 |
| GO:0072596 | establishment of protein localization to chloroplast | 4.95E-07 | 0.000126 |
| GO:0072598 | protein localization to chloroplast | 4.95E-07 | 0.000126 |
| GO:0045036 | protein targeting to chloroplast | 4.95E-07 | 0.000126 |
| GO:0006379 | mRNA cleavage | 5.16E-07 | 0.000126 |
| GO:0006397 | mRNA processing | 5.61E-07 | 0.000126 |
| GO:0008033 | tRNA processing | 5.84E-07 | 0.000126 |
| GO:0016441 | posttranscriptional gene silencing | 6.28E-07 | 0.000126 |
| GO:0006446 | regulation of translational initiation | 6.38E-07 | 0.000126 |
| GO:0045038 | protein import into chloroplast thylakoid membrane | 6.49E-07 | 0.000126 |
| GO:0010608 | posttranscriptional regulation of gene expression | 9.33E-07 | 0.000172 |
| GO:0034470 | ncRNA processing | 1.11E-06 | 0.000184 |
| GO:0017038 | protein import | 1.15E-06 | 0.000184 |
| GO:0071806 | protein transmembrane transport | 1.22E-06 | 0.000184 |
| GO:0065002 | intracellular protein transmembrane transport | 1.22E-06 | 0.000184 |
| GO:0006418 | tRNA aminoacylation for protein translation | 1.52E-06 | 0.000201 |
| GO:0043039 | tRNA aminoacylation | 1.52E-06 | 0.000201 |
| GO:0043038 | amino acid activation | 1.52E-06 | 0.000201 |
| GO:0016071 | mRNA metabolic process | 1.75E-06 | 0.000223 |
| GO:0006281 | DNA repair | 1.85E-06 | 0.000227 |
| GO:0006399 | tRNA metabolic process | 2.01E-06 | 0.000233 |
| GO:0006974 | cellular response to DNA damage stimulus | 2.26E-06 | 0.000235 |
| GO:0006396 | RNA processing | 2.28E-06 | 0.000235 |
| GO:0034660 | ncRNA metabolic process | 2.34E-06 | 0.000235 |
| GO:0006413 | translational initiation | 3.07E-06 | 0.000291 |
| GO:0051716 | cellular response to stimulus | 3.08E-06 | 0.000291 |
| GO:0043933 | protein-containing complex subunit organization | 3.19E-06 | 0.000293 |
| GO:0009793 | embryo development ending in seed dormancy | 3.52E-06 | 0.00031 |
| GO:0006259 | DNA metabolic process | 3.62E-06 | 0.00031 |
| GO:0033554 | cellular response to stress | 3.69E-06 | 0.00031 |
| GO:0009790 | embryo development | 4.03E-06 | 0.000318 |
| GO:0006325 | chromatin organization | 4.3E-06 | 0.000318 |
| GO:0034622 | cellular protein-containing complex assembly | 4.38E-06 | 0.000318 |
| GO:0007275 | multicellular organism development | 4.41E-06 | 0.000318 |
| GO:0016070 | RNA metabolic process | 4.55E-06 | 0.000321 |
| GO:0090304 | nucleic acid metabolic process | 4.82E-06 | 0.000326 |
| GO:0006139 | nucleobase-containing compound metabolic process | 5.81E-06 | 0.000348 |
| GO:0051603 | proteolysis involved in cellular protein catabolic process | 5.93E-06 | 0.000348 |
| GO:0046907 | intracellular transport | 6.06E-06 | 0.000348 |
| GO:0003006 | developmental process involved in reproduction | 6.13E-06 | 0.000348 |
| GO:0022414 | reproductive process | 6.35E-06 | 0.000348 |
| GO:0048856 | anatomical structure development | 6.59E-06 | 0.000348 |
| GO:0032501 | multicellular organismal process | 6.75E-06 | 0.000348 |
| GO:0046483 | heterocycle metabolic process | 6.84E-06 | 0.000348 |
| GO:0006419 | alanyl-tRNA aminoacylation | 6.86E-06 | 0.000348 |
| GO:0006725 | cellular aromatic compound metabolic process | 6.94E-06 | 0.000348 |
| GO:0017145 | stem cell division | 7.17E-06 | 0.000348 |
| GO:0010224 | response to UV-B | 7.33E-06 | 0.000348 |
| GO:1901360 | organic cyclic compound metabolic process | 7.36E-06 | 0.000348 |
| GO:0034641 | cellular nitrogen compound metabolic process | 7.69E-06 | 0.000348 |
| GO:0048519 | negative regulation of biological process | 8.3E-06 | 0.000348 |
| GO:0000377 | RNA splicing, via transesterification reactions with bulged adenosine as nucleophile | 8.33E-06 | 0.000348 |
| GO:0000375 | RNA splicing, via transesterification reactions | 8.33E-06 | 0.000348 |
| GO:0044249 | cellular biosynthetic process | 8.61E-06 | 0.000348 |
| GO:0043632 | modification-dependent macromolecule catabolic process | 9.26E-06 | 0.000357 |
| GO:0006511 | ubiquitin-dependent protein catabolic process | 9.26E-06 | 0.000357 |
| GO:0019941 | modification-dependent protein catabolic process | 9.26E-06 | 0.000357 |
| GO:0006807 | nitrogen compound metabolic process | 1.08E-05 | 0.000403 |
| GO:0043170 | macromolecule metabolic process | 1.09E-05 | 0.000403 |
| GO:0065003 | protein-containing complex assembly | 1.1E-05 | 0.000403 |
| GO:0032774 | RNA biosynthetic process | 1.14E-05 | 0.000416 |
| GO:0044237 | cellular metabolic process | 1.22E-05 | 0.00044 |
| GO:0051649 | establishment of localization in cell | 1.27E-05 | 0.000451 |
| GO:0050657 | nucleic acid transport | 1.43E-05 | 0.000489 |
| GO:0006405 | RNA export from nucleus | 1.43E-05 | 0.000489 |
| GO:0051236 | establishment of RNA localization | 1.43E-05 | 0.000489 |
| GO:0050658 | RNA transport | 1.43E-05 | 0.000489 |
| GO:0051641 | cellular localization | 1.47E-05 | 0.000491 |
| GO:0033365 | protein localization to organelle | 1.48E-05 | 0.000491 |
| GO:0009987 | cellular process | 1.51E-05 | 0.000494 |
| GO:0044265 | cellular macromolecule catabolic process | 1.86E-05 | 0.000596 |
| GO:0032502 | developmental process | 1.9E-05 | 0.000605 |
| GO:0000373 | Group II intron splicing | 1.99E-05 | 0.000628 |
| GO:0008380 | RNA splicing | 2.34E-05 | 0.000731 |
| GO:0034654 | nucleobase-containing compound biosynthetic process | 2.44E-05 | 0.000756 |
| GO:0044271 | cellular nitrogen compound biosynthetic process | 2.65E-05 | 0.000806 |
| GO:0006520 | cellular amino acid metabolic process | 2.65E-05 | 0.000806 |
| GO:0044238 | primary metabolic process | 2.74E-05 | 0.000826 |
| GO:0098656 | anion transmembrane transport | 2.78E-05 | 0.00083 |
| GO:0010038 | response to metal ion | 3.42E-05 | 0.001012 |
| GO:0016458 | gene silencing | 3.71E-05 | 0.001087 |
| GO:1901576 | organic substance biosynthetic process | 3.84E-05 | 0.001097 |
| GO:0009756 | carbohydrate mediated signaling | 3.84E-05 | 0.001097 |
| GO:0010182 | sugar mediated signaling pathway | 3.84E-05 | 0.001097 |
| GO:0010605 | negative regulation of macromolecule metabolic process | 4.77E-05 | 0.001295 |
| GO:0031119 | tRNA pseudouridine synthesis | 4.77E-05 | 0.001295 |
| GO:0006352 | DNA-templated transcription, initiation | 4.78E-05 | 0.001295 |
| GO:0009127 | purine nucleoside monophosphate biosynthetic process | 4.81E-05 | 0.001295 |
| GO:0009168 | purine ribonucleoside monophosphate biosynthetic process | 4.81E-05 | 0.001295 |
| GO:0009167 | purine ribonucleoside monophosphate metabolic process | 4.81E-05 | 0.001295 |
| GO:0009126 | purine nucleoside monophosphate metabolic process | 4.81E-05 | 0.001295 |
| GO:0006886 | intracellular protein transport | 4.87E-05 | 0.001301 |
| GO:0009058 | biosynthetic process | 5.43E-05 | 0.001433 |
| GO:0000725 | recombinational repair | 5.51E-05 | 0.001433 |
| GO:0000724 | double-strand break repair via homologous recombination | 5.51E-05 | 0.001433 |
| GO:0051028 | mRNA transport | 5.58E-05 | 0.001433 |
| GO:0006406 | mRNA export from nucleus | 5.58E-05 | 0.001433 |
| GO:0008152 | metabolic process | 5.94E-05 | 0.001513 |
| GO:0009059 | macromolecule biosynthetic process | 6.53E-05 | 0.001651 |
| GO:0046686 | response to cadmium ion | 7.13E-05 | 0.001776 |
| GO:0043487 | regulation of RNA stability | 7.86E-05 | 0.001794 |
| GO:0043488 | regulation of mRNA stability | 7.86E-05 | 0.001794 |
| GO:0048255 | mRNA stabilization | 7.86E-05 | 0.001794 |
| GO:1903312 | negative regulation of mRNA metabolic process | 7.86E-05 | 0.001794 |
| GO:0009895 | negative regulation of catabolic process | 7.86E-05 | 0.001794 |
| GO:0043489 | RNA stabilization | 7.86E-05 | 0.001794 |
| GO:0031330 | negative regulation of cellular catabolic process | 7.86E-05 | 0.001794 |
| GO:1902369 | negative regulation of RNA catabolic process | 7.86E-05 | 0.001794 |
| GO:1902373 | negative regulation of mRNA catabolic process | 7.86E-05 | 0.001794 |
| GO:1903311 | regulation of mRNA metabolic process | 7.86E-05 | 0.001794 |
| GO:0061013 | regulation of mRNA catabolic process | 7.86E-05 | 0.001794 |
| GO:0071704 | organic substance metabolic process | 8.24E-05 | 0.001868 |
| GO:0034613 | cellular protein localization | 8.35E-05 | 0.001876 |
| GO:0009893 | positive regulation of metabolic process | 8.39E-05 | 0.001876 |
| GO:0031123 | RNA 3'-end processing | 8.88E-05 | 0.001974 |
| GO:0010035 | response to inorganic substance | 9.19E-05 | 0.002028 |
| GO:0046506 | sulfolipid biosynthetic process | 9.37E-05 | 0.002041 |
| GO:0046505 | sulfolipid metabolic process | 9.37E-05 | 0.002041 |
| GO:0006809 | nitric oxide biosynthetic process | 9.71E-05 | 0.002089 |
| GO:0046209 | nitric oxide metabolic process | 9.71E-05 | 0.002089 |
| GO:1901463 | regulation of tetrapyrrole biosynthetic process | 0.000102 | 0.002168 |
| GO:0010380 | regulation of chlorophyll biosynthetic process | 0.000102 | 0.002168 |
| GO:0006298 | mismatch repair | 0.000109 | 0.00229 |
| GO:0006302 | double-strand break repair | 0.00011 | 0.002313 |
| GO:0006612 | protein targeting to membrane | 0.000116 | 0.002413 |
| GO:0032984 | protein-containing complex disassembly | 0.000119 | 0.002469 |
| GO:0031124 | mRNA 3'-end processing | 0.000128 | 0.002607 |
| GO:0006378 | mRNA polyadenylation | 0.000128 | 0.002607 |
| GO:0072594 | establishment of protein localization to organelle | 0.00013 | 0.002638 |
| GO:0051168 | nuclear export | 0.000133 | 0.002684 |
| GO:0010628 | positive regulation of gene expression | 0.000142 | 0.002842 |
| GO:0009057 | macromolecule catabolic process | 0.000146 | 0.002887 |
| GO:0009102 | biotin biosynthetic process | 0.000168 | 0.003268 |
| GO:0006768 | biotin metabolic process | 0.000168 | 0.003268 |
| GO:0009411 | response to UV | 0.000177 | 0.003422 |
| GO:0000338 | protein deneddylation | 0.000187 | 0.003601 |
| GO:0009892 | negative regulation of metabolic process | 0.000207 | 0.003941 |
| GO:0046351 | disaccharide biosynthetic process | 0.000207 | 0.003941 |
| GO:0045005 | DNA-dependent DNA replication maintenance of fidelity | 0.000218 | 0.004132 |
| GO:0090056 | regulation of chlorophyll metabolic process | 0.000224 | 0.004199 |
| GO:1901401 | regulation of tetrapyrrole metabolic process | 0.000224 | 0.004199 |
| GO:0010387 | COP9 signalosome assembly | 0.000229 | 0.004227 |
| GO:0046033 | AMP metabolic process | 0.000232 | 0.004242 |
| GO:0006167 | AMP biosynthetic process | 0.000232 | 0.004242 |
| GO:0010604 | positive regulation of macromolecule metabolic process | 0.000264 | 0.004777 |
| GO:0009658 | chloroplast organization | 0.000264 | 0.004777 |
| GO:0046037 | GMP metabolic process | 0.000288 | 0.005152 |
| GO:0006177 | GMP biosynthetic process | 0.000288 | 0.005152 |
| GO:0016043 | cellular component organization | 0.000306 | 0.005418 |
| GO:0006415 | translational termination | 0.000311 | 0.005418 |
| GO:0043624 | cellular protein complex disassembly | 0.000311 | 0.005418 |
| GO:0009639 | response to red or far red light | 0.000322 | 0.005578 |
| GO:0006082 | organic acid metabolic process | 0.000332 | 0.005729 |
| GO:0006605 | protein targeting | 0.000423 | 0.007222 |
| GO:0070646 | protein modification by small protein removal | 0.000458 | 0.007777 |
| GO:0070727 | cellular macromolecule localization | 0.000463 | 0.007787 |
| GO:0044281 | small molecule metabolic process | 0.000474 | 0.007885 |
| GO:0006414 | translational elongation | 0.000477 | 0.007904 |
| GO:0009156 | ribonucleoside monophosphate biosynthetic process | 0.000494 | 0.008095 |
| GO:0009124 | nucleoside monophosphate biosynthetic process | 0.000494 | 0.008095 |
| GO:1902531 | regulation of intracellular signal transduction | 0.000534 | 0.008663 |
| GO:0006338 | chromatin remodeling | 0.000548 | 0.008776 |
| GO:0051169 | nuclear transport | 0.000549 | 0.008776 |
| GO:0006913 | nucleocytoplasmic transport | 0.000549 | 0.008776 |
| GO:0022607 | cellular component assembly | 0.000559 | 0.008895 |
| GO:0009161 | ribonucleoside monophosphate metabolic process | 0.00059 | 0.009307 |
| GO:0009123 | nucleoside monophosphate metabolic process | 0.00059 | 0.009307 |

**Supplemental Table 5. Enriched GO terms upregulated in only roots under desiccation.**

| **GO** | **GO term description** | **p_uncorrected** | | **p_fdr_bh** |
| --- | --- | --- | --- | --- |
| GO:0046677 | response to antibiotic | | 2.19E-07 | 0.000137 |
| GO:0060341 | regulation of cellular localization | | 2.48E-07 | 0.000137 |
| GO:0030581 | symbiont intracellular protein transport in host | | 3.02E-07 | 0.000137 |
| GO:0043562 | cellular response to nitrogen levels | | 3.38E-07 | 0.000137 |
| GO:0042744 | hydrogen peroxide catabolic process | | 3.46E-07 | 0.000137 |
| GO:0010099 | regulation of photomorphogenesis | | 6.2E-07 | 0.000137 |
| GO:2000030 | regulation of response to red or far red light | | 6.2E-07 | 0.000137 |
| GO:0042542 | response to hydrogen peroxide | | 7.34E-07 | 0.000137 |
| GO:0015986 | ATP synthesis coupled proton transport | | 7.53E-07 | 0.000137 |
| GO:0015985 | energy coupled proton transport, down electrochemical gradient | | 7.53E-07 | 0.000137 |
| GO:0006995 | cellular response to nitrogen starvation | | 7.59E-07 | 0.000137 |
| GO:0009970 | cellular response to sulfate starvation | | 7.59E-07 | 0.000137 |
| GO:0009062 | fatty acid catabolic process | | 7.82E-07 | 0.000137 |
| GO:0016050 | vesicle organization | | 8.03E-07 | 0.000137 |
| GO:0006635 | fatty acid beta-oxidation | | 8.24E-07 | 0.000137 |
| GO:0015800 | acidic amino acid transport | | 8.75E-07 | 0.000137 |
| GO:0031163 | metallo-sulfur cluster assembly | | 8.85E-07 | 0.000137 |
| GO:0016226 | iron-sulfur cluster assembly | | 8.85E-07 | 0.000137 |
| GO:0006891 | intra-Golgi vesicle-mediated transport | | 9.19E-07 | 0.000137 |
| GO:0015711 | organic anion transport | | 9.76E-07 | 0.000137 |
| GO:0048227 | plasma membrane to endosome transport | | 1.03E-06 | 0.000137 |
| GO:0010117 | photoprotection | | 1.09E-06 | 0.000137 |
| GO:0072593 | reactive oxygen species metabolic process | | 1.22E-06 | 0.000137 |
| GO:0009853 | photorespiration | | 1.37E-06 | 0.000137 |
| GO:0009063 | cellular amino acid catabolic process | | 1.41E-06 | 0.000137 |
| GO:0006754 | ATP biosynthetic process | | 1.41E-06 | 0.000137 |
| GO:0016311 | dephosphorylation | | 1.43E-06 | 0.000137 |
| GO:0034220 | ion transmembrane transport | | 1.54E-06 | 0.000137 |
| GO:0046395 | carboxylic acid catabolic process | | 1.61E-06 | 0.000137 |
| GO:0006470 | protein dephosphorylation | | 1.64E-06 | 0.000137 |
| GO:0030258 | lipid modification | | 1.67E-06 | 0.000137 |
| GO:0031539 | positive regulation of anthocyanin metabolic process | | 1.8E-06 | 0.000137 |
| GO:0000302 | response to reactive oxygen species | | 1.82E-06 | 0.000137 |
| GO:0016054 | organic acid catabolic process | | 1.84E-06 | 0.000137 |
| GO:1901606 | alpha-amino acid catabolic process | | 1.86E-06 | 0.000137 |
| GO:0009644 | response to high light intensity | | 2E-06 | 0.000137 |
| GO:1902600 | proton transmembrane transport | | 2.05E-06 | 0.000137 |
| GO:0006612 | protein targeting to membrane | | 2.05E-06 | 0.000137 |
| GO:0061024 | membrane organization | | 2.1E-06 | 0.000137 |
| GO:0044282 | small molecule catabolic process | | 2.11E-06 | 0.000137 |
| GO:0009642 | response to light intensity | | 2.19E-06 | 0.000137 |
| GO:0048193 | Golgi vesicle transport | | 2.19E-06 | 0.000137 |
| GO:0006839 | mitochondrial transport | | 2.19E-06 | 0.000137 |
| GO:0009142 | nucleoside triphosphate biosynthetic process | | 2.2E-06 | 0.000137 |
| GO:0061025 | membrane fusion | | 2.22E-06 | 0.000137 |
| GO:1901565 | organonitrogen compound catabolic process | | 2.26E-06 | 0.000137 |
| GO:0098660 | inorganic ion transmembrane transport | | 2.35E-06 | 0.000137 |
| GO:0098655 | cation transmembrane transport | | 2.35E-06 | 0.000137 |
| GO:0098662 | inorganic cation transmembrane transport | | 2.35E-06 | 0.000137 |
| GO:0030163 | protein catabolic process | | 2.36E-06 | 0.000137 |
| GO:0043094 | cellular metabolic compound salvage | | 2.43E-06 | 0.000137 |
| GO:0009144 | purine nucleoside triphosphate metabolic process | | 2.63E-06 | 0.000137 |
| GO:0009205 | purine ribonucleoside triphosphate metabolic process | | 2.63E-06 | 0.000137 |
| GO:0009206 | purine ribonucleoside triphosphate biosynthetic process | | 2.63E-06 | 0.000137 |
| GO:0009201 | ribonucleoside triphosphate biosynthetic process | | 2.63E-06 | 0.000137 |
| GO:0009199 | ribonucleoside triphosphate metabolic process | | 2.63E-06 | 0.000137 |
| GO:0009145 | purine nucleoside triphosphate biosynthetic process | | 2.63E-06 | 0.000137 |
| GO:0010353 | response to trehalose | | 2.64E-06 | 0.000137 |
| GO:0015904 | tetracycline transmembrane transport | | 2.64E-06 | 0.000137 |
| GO:0019395 | fatty acid oxidation | | 2.67E-06 | 0.000137 |
| GO:0034440 | lipid oxidation | | 2.67E-06 | 0.000137 |
| GO:0042816 | vitamin B6 metabolic process | | 2.91E-06 | 0.000137 |
| GO:0042819 | vitamin B6 biosynthetic process | | 2.91E-06 | 0.000137 |
| GO:0008614 | pyridoxine metabolic process | | 2.91E-06 | 0.000137 |
| GO:0008615 | pyridoxine biosynthetic process | | 2.91E-06 | 0.000137 |
| GO:0009141 | nucleoside triphosphate metabolic process | | 2.92E-06 | 0.000137 |
| GO:0046034 | ATP metabolic process | | 2.96E-06 | 0.000137 |
| GO:0071496 | cellular response to external stimulus | | 2.98E-06 | 0.000137 |
| GO:0031668 | cellular response to extracellular stimulus | | 2.98E-06 | 0.000137 |
| GO:0051603 | proteolysis involved in cellular protein catabolic process | | 3.09E-06 | 0.00014 |
| GO:0016192 | vesicle-mediated transport | | 3.33E-06 | 0.000144 |
| GO:0009408 | response to heat | | 3.46E-06 | 0.000144 |
| GO:0009991 | response to extracellular stimulus | | 3.47E-06 | 0.000144 |
| GO:0009057 | macromolecule catabolic process | | 3.79E-06 | 0.000149 |
| GO:0006790 | sulfur compound metabolic process | | 3.98E-06 | 0.000151 |
| GO:0006979 | response to oxidative stress | | 4.03E-06 | 0.000151 |
| GO:0044265 | cellular macromolecule catabolic process | | 4.04E-06 | 0.000151 |
| GO:0019941 | modification-dependent protein catabolic process | | 4.15E-06 | 0.000151 |
| GO:0043632 | modification-dependent macromolecule catabolic process | | 4.15E-06 | 0.000151 |
| GO:0006511 | ubiquitin-dependent protein catabolic process | | 4.15E-06 | 0.000151 |
| GO:0006099 | tricarboxylic acid cycle | | 4.27E-06 | 0.000153 |
| GO:0009737 | response to abscisic acid | | 4.41E-06 | 0.000153 |
| GO:0006886 | intracellular protein transport | | 4.45E-06 | 0.000153 |
| GO:0045184 | establishment of protein localization | | 4.53E-06 | 0.000154 |
| GO:0015833 | peptide transport | | 4.56E-06 | 0.000154 |
| GO:0008104 | protein localization | | 4.69E-06 | 0.000156 |
| GO:0097305 | response to alcohol | | 4.7E-06 | 0.000156 |
| GO:0046686 | response to cadmium ion | | 4.86E-06 | 0.000157 |
| GO:0042886 | amide transport | | 4.9E-06 | 0.000157 |
| GO:0071705 | nitrogen compound transport | | 4.92E-06 | 0.000157 |
| GO:0046907 | intracellular transport | | 4.96E-06 | 0.000157 |
| GO:0051641 | cellular localization | | 4.99E-06 | 0.000157 |
| GO:0051649 | establishment of localization in cell | | 5.08E-06 | 0.000157 |
| GO:0033036 | macromolecule localization | | 5.09E-06 | 0.000157 |
| GO:0010038 | response to metal ion | | 5.31E-06 | 0.00016 |
| GO:0046487 | glyoxylate metabolic process | | 5.33E-06 | 0.00016 |
| GO:0015031 | protein transport | | 5.36E-06 | 0.00016 |
| GO:1900180 | regulation of protein localization to nucleus | | 5.57E-06 | 0.000161 |
| GO:0042306 | regulation of protein import into nucleus | | 5.57E-06 | 0.000161 |
| GO:1904589 | regulation of protein import | | 5.57E-06 | 0.000161 |
| GO:0006970 | response to osmotic stress | | 5.58E-06 | 0.000161 |
| GO:0044248 | cellular catabolic process | | 5.65E-06 | 0.000161 |
| GO:0009266 | response to temperature stimulus | | 5.77E-06 | 0.000161 |
| GO:0009651 | response to salt stress | | 5.8E-06 | 0.000161 |
| GO:0009056 | catabolic process | | 6.15E-06 | 0.000161 |
| GO:0072329 | monocarboxylic acid catabolic process | | 6.28E-06 | 0.000161 |
| GO:0019430 | removal of superoxide radicals | | 6.33E-06 | 0.000161 |
| GO:0098869 | cellular oxidant detoxification | | 6.33E-06 | 0.000161 |
| GO:1990748 | cellular detoxification | | 6.33E-06 | 0.000161 |
| GO:1901575 | organic substance catabolic process | | 6.37E-06 | 0.000161 |
| GO:0070972 | protein localization to endoplasmic reticulum | | 6.37E-06 | 0.000161 |
| GO:0006613 | cotranslational protein targeting to membrane | | 6.37E-06 | 0.000161 |
| GO:0006614 | SRP-dependent cotranslational protein targeting to membrane | | 6.37E-06 | 0.000161 |
| GO:0072599 | establishment of protein localization to endoplasmic reticulum | | 6.37E-06 | 0.000161 |
| GO:0045047 | protein targeting to ER | | 6.37E-06 | 0.000161 |
| GO:0010035 | response to inorganic substance | | 6.48E-06 | 0.000162 |
| GO:0071702 | organic substance transport | | 6.73E-06 | 0.000166 |
| GO:0051336 | regulation of hydrolase activity | | 6.91E-06 | 0.000167 |
| GO:0006820 | anion transport | | 7.06E-06 | 0.000167 |
| GO:1901700 | response to oxygen-containing compound | | 7.22E-06 | 0.000167 |
| GO:1903827 | regulation of cellular protein localization | | 7.38E-06 | 0.000167 |
| GO:0070201 | regulation of establishment of protein localization | | 7.38E-06 | 0.000167 |
| GO:0051223 | regulation of protein transport | | 7.38E-06 | 0.000167 |
| GO:0033157 | regulation of intracellular protein transport | | 7.38E-06 | 0.000167 |
| GO:0090087 | regulation of peptide transport | | 7.38E-06 | 0.000167 |
| GO:0046822 | regulation of nucleocytoplasmic transport | | 7.38E-06 | 0.000167 |
| GO:0032386 | regulation of intracellular transport | | 7.38E-06 | 0.000167 |
| GO:0006811 | ion transport | | 7.47E-06 | 0.000167 |
| GO:0072524 | pyridine-containing compound metabolic process | | 7.8E-06 | 0.000171 |
| GO:0072525 | pyridine-containing compound biosynthetic process | | 7.8E-06 | 0.000171 |
| GO:0007034 | vacuolar transport | | 8.12E-06 | 0.000177 |
| GO:1901698 | response to nitrogen compound | | 8.87E-06 | 0.000191 |
| GO:0097437 | maintenance of dormancy | | 9.22E-06 | 0.000193 |
| GO:0010231 | maintenance of seed dormancy | | 9.22E-06 | 0.000193 |
| GO:0009628 | response to abiotic stimulus | | 9.41E-06 | 0.000194 |
| GO:0006810 | transport | | 9.68E-06 | 0.000194 |
| GO:0042221 | response to chemical | | 9.71E-06 | 0.000194 |
| GO:0051179 | localization | | 9.84E-06 | 0.000194 |
| GO:0042594 | response to starvation | | 9.87E-06 | 0.000194 |
| GO:0051234 | establishment of localization | | 1.03E-05 | 0.000197 |
| GO:0006081 | cellular aldehyde metabolic process | | 1.03E-05 | 0.000197 |
| GO:0006950 | response to stress | | 1.11E-05 | 0.000203 |
| GO:0031667 | response to nutrient levels | | 1.12E-05 | 0.000203 |
| GO:0006605 | protein targeting | | 1.22E-05 | 0.000219 |
| GO:0050896 | response to stimulus | | 1.46E-05 | 0.000253 |
| GO:0072657 | protein localization to membrane | | 1.48E-05 | 0.000254 |
| GO:0090150 | establishment of protein localization to membrane | | 1.48E-05 | 0.000254 |
| GO:0009416 | response to light stimulus | | 1.5E-05 | 0.000256 |
| GO:0031669 | cellular response to nutrient levels | | 1.61E-05 | 0.000273 |
| GO:0044247 | cellular polysaccharide catabolic process | | 1.67E-05 | 0.00028 |
| GO:0009251 | glucan catabolic process | | 1.67E-05 | 0.00028 |
| GO:0034613 | cellular protein localization | | 1.75E-05 | 0.000292 |
| GO:0009150 | purine ribonucleotide metabolic process | | 1.87E-05 | 0.000306 |
| GO:0009165 | nucleotide biosynthetic process | | 1.89E-05 | 0.000306 |
| GO:0010205 | photoinhibition | | 1.9E-05 | 0.000306 |
| GO:0043155 | negative regulation of photosynthesis, light reaction | | 1.9E-05 | 0.000306 |
| GO:1905156 | negative regulation of photosynthesis | | 1.9E-05 | 0.000306 |
| GO:0046339 | diacylglycerol metabolic process | | 1.9E-05 | 0.000306 |
| GO:0006651 | diacylglycerol biosynthetic process | | 1.9E-05 | 0.000306 |
| GO:0009987 | cellular process | | 1.97E-05 | 0.000316 |
| GO:0005983 | starch catabolic process | | 2.01E-05 | 0.00032 |
| GO:0042743 | hydrogen peroxide metabolic process | | 2.04E-05 | 0.000324 |
| GO:0006163 | purine nucleotide metabolic process | | 2.13E-05 | 0.00033 |
| GO:0006623 | protein targeting to vacuole | | 2.13E-05 | 0.00033 |
| GO:0072666 | establishment of protein localization to vacuole | | 2.13E-05 | 0.00033 |
| GO:0072665 | protein localization to vacuole | | 2.13E-05 | 0.00033 |
| GO:0042548 | regulation of photosynthesis, light reaction | | 2.15E-05 | 0.000331 |
| GO:0006801 | superoxide metabolic process | | 2.24E-05 | 0.000344 |
| GO:0009259 | ribonucleotide metabolic process | | 2.31E-05 | 0.000352 |
| GO:0009648 | photoperiodism | | 2.34E-05 | 0.000354 |
| GO:0008285 | negative regulation of cell population proliferation | | 2.34E-05 | 0.000354 |
| GO:0010227 | floral organ abscission | | 2.45E-05 | 0.000367 |
| GO:0009838 | abscission | | 2.45E-05 | 0.000367 |
| GO:0072521 | purine-containing compound metabolic process | | 3.18E-05 | 0.00047 |
| GO:0009314 | response to radiation | | 3.22E-05 | 0.000473 |
| GO:0019693 | ribose phosphate metabolic process | | 3.36E-05 | 0.000492 |
| GO:0009267 | cellular response to starvation | | 3.41E-05 | 0.000498 |
| GO:0006753 | nucleoside phosphate metabolic process | | 3.89E-05 | 0.000564 |
| GO:0010243 | response to organonitrogen compound | | 4.15E-05 | 0.0006 |
| GO:0016036 | cellular response to phosphate starvation | | 4.35E-05 | 0.000626 |
| GO:0009260 | ribonucleotide biosynthetic process | | 4.45E-05 | 0.000638 |
| GO:0043087 | regulation of GTPase activity | | 4.48E-05 | 0.000639 |
| GO:0034311 | diol metabolic process | | 5.06E-05 | 0.000716 |
| GO:0034312 | diol biosynthetic process | | 5.06E-05 | 0.000716 |
| GO:1901293 | nucleoside phosphate biosynthetic process | | 5.15E-05 | 0.000725 |
| GO:0008380 | RNA splicing | | 5.17E-05 | 0.000725 |
| GO:0009110 | vitamin biosynthetic process | | 5.66E-05 | 0.000781 |
| GO:0016197 | endosomal transport | | 5.92E-05 | 0.00081 |
| GO:0006457 | protein folding | | 6.37E-05 | 0.000864 |
| GO:0032928 | regulation of superoxide anion generation | | 6.53E-05 | 0.000876 |
| GO:0090322 | regulation of superoxide metabolic process | | 6.53E-05 | 0.000876 |
| GO:0015672 | monovalent inorganic cation transport | | 6.77E-05 | 0.000904 |
| GO:0009117 | nucleotide metabolic process | | 6.85E-05 | 0.000909 |
| GO:0010109 | regulation of photosynthesis | | 7.36E-05 | 0.000967 |
| GO:0046390 | ribose phosphate biosynthetic process | | 7.39E-05 | 0.000967 |
| GO:0006842 | tricarboxylic acid transport | | 7.78E-05 | 0.001006 |
| GO:0015746 | citrate transport | | 7.78E-05 | 0.001006 |
| GO:0032880 | regulation of protein localization | | 8E-05 | 0.001031 |
| GO:0048523 | negative regulation of cellular process | | 8.12E-05 | 0.001042 |
| GO:0009152 | purine ribonucleotide biosynthetic process | | 8.38E-05 | 0.001071 |
| GO:0042817 | pyridoxal metabolic process | | 8.41E-05 | 0.001071 |
| GO:0006766 | vitamin metabolic process | | 9.63E-05 | 0.001203 |
| GO:0009939 | positive regulation of gibberellic acid mediated signaling pathway | | 9.95E-05 | 0.001234 |
| GO:0006164 | purine nucleotide biosynthetic process | | 0.000101 | 0.001239 |
| GO:0018130 | heterocycle biosynthetic process | | 0.000101 | 0.001242 |
| GO:0019637 | organophosphate metabolic process | | 0.000122 | 0.001478 |
| GO:0044242 | cellular lipid catabolic process | | 0.000124 | 0.001493 |
| GO:0010200 | response to chitin | | 0.000134 | 0.001588 |
| GO:0000377 | RNA splicing, via transesterification reactions with bulged adenosine as nucleophile | | 0.000134 | 0.001588 |
| GO:0000375 | RNA splicing, via transesterification reactions | | 0.000134 | 0.001588 |
| GO:0022904 | respiratory electron transport chain | | 0.000134 | 0.001588 |
| GO:0032879 | regulation of localization | | 0.000137 | 0.001616 |
| GO:0046471 | phosphatidylglycerol metabolic process | | 0.000165 | 0.001909 |
| GO:0010335 | response to non-ionic osmotic stress | | 0.000165 | 0.001909 |
| GO:0006982 | response to lipid hydroperoxide | | 0.000165 | 0.001909 |
| GO:0006655 | phosphatidylglycerol biosynthetic process | | 0.000165 | 0.001909 |
| GO:0033194 | response to hydroperoxide | | 0.000165 | 0.001909 |
| GO:0090407 | organophosphate biosynthetic process | | 0.000186 | 0.002131 |
| GO:0043289 | apocarotenoid biosynthetic process | | 0.000194 | 0.002195 |
| GO:0009688 | abscisic acid biosynthetic process | | 0.000194 | 0.002195 |
| GO:1902645 | tertiary alcohol biosynthetic process | | 0.000194 | 0.002195 |
| GO:0043269 | regulation of ion transport | | 0.000197 | 0.002227 |
| GO:0031324 | negative regulation of cellular metabolic process | | 0.0002 | 0.002254 |
| GO:0045454 | cell redox homeostasis | | 0.000205 | 0.0023 |
| GO:0006896 | Golgi to vacuole transport | | 0.000209 | 0.002327 |
| GO:0006892 | post-Golgi vesicle-mediated transport | | 0.000209 | 0.002327 |
| GO:0007041 | lysosomal transport | | 0.000212 | 0.002327 |
| GO:0009173 | pyrimidine ribonucleoside monophosphate metabolic process | | 0.000212 | 0.002327 |
| GO:0009129 | pyrimidine nucleoside monophosphate metabolic process | | 0.000212 | 0.002327 |
| GO:0008333 | endosome to lysosome transport | | 0.000212 | 0.002327 |
| GO:0072594 | establishment of protein localization to organelle | | 0.000229 | 0.002504 |
| GO:0048194 | Golgi vesicle budding | | 0.000232 | 0.002518 |
| GO:0000272 | polysaccharide catabolic process | | 0.000232 | 0.002518 |
| GO:0070727 | cellular macromolecule localization | | 0.000233 | 0.002518 |
| GO:0018131 | oxazole or thiazole biosynthetic process | | 0.000273 | 0.002929 |
| GO:0046484 | oxazole or thiazole metabolic process | | 0.000273 | 0.002929 |
| GO:0006097 | glyoxylate cycle | | 0.000277 | 0.002962 |
| GO:0042538 | hyperosmotic salinity response | | 0.000288 | 0.003064 |
| GO:0033523 | histone H2B ubiquitination | | 0.000291 | 0.003065 |
| GO:0016574 | histone ubiquitination | | 0.000291 | 0.003065 |
| GO:0055086 | nucleobase-containing small molecule metabolic process | | 0.000299 | 0.003137 |
| GO:0043085 | positive regulation of catalytic activity | | 0.000307 | 0.003183 |
| GO:0044093 | positive regulation of molecular function | | 0.000307 | 0.003183 |
| GO:0000398 | mRNA splicing, via spliceosome | | 0.000311 | 0.003215 |
| GO:0050826 | response to freezing | | 0.000313 | 0.003227 |
| GO:0010039 | response to iron ion | | 0.000314 | 0.003227 |
| GO:0031365 | N-terminal protein amino acid modification | | 0.000334 | 0.003405 |
| GO:0006499 | N-terminal protein myristoylation | | 0.000335 | 0.003405 |
| GO:0018377 | protein myristoylation | | 0.000335 | 0.003405 |
| GO:0006498 | N-terminal protein lipidation | | 0.000335 | 0.003405 |
| GO:0043543 | protein acylation | | 0.000373 | 0.003775 |
| GO:0022611 | dormancy process | | 0.000375 | 0.003784 |
| GO:0009409 | response to cold | | 0.000399 | 0.003996 |
| GO:0043467 | regulation of generation of precursor metabolites and energy | | 0.000401 | 0.003998 |
| GO:0007030 | Golgi organization | | 0.000403 | 0.004005 |
| GO:0042364 | water-soluble vitamin biosynthetic process | | 0.000406 | 0.004025 |
| GO:0019458 | methionine catabolic process via 2-oxobutanoate | | 0.000414 | 0.004029 |
| GO:0006562 | proline catabolic process | | 0.000414 | 0.004029 |
| GO:0009087 | methionine catabolic process | | 0.000414 | 0.004029 |
| GO:0000098 | sulfur amino acid catabolic process | | 0.000414 | 0.004029 |
| GO:0007154 | cell communication | | 0.000419 | 0.004072 |
| GO:0035264 | multicellular organism growth | | 0.000472 | 0.004514 |
| GO:0044237 | cellular metabolic process | | 0.000475 | 0.00453 |
| GO:0016108 | tetraterpenoid metabolic process | | 0.000508 | 0.004768 |
| GO:0016116 | carotenoid metabolic process | | 0.000508 | 0.004768 |
| GO:0033365 | protein localization to organelle | | 0.00051 | 0.004768 |
| GO:0072522 | purine-containing compound biosynthetic process | | 0.000538 | 0.005016 |
| GO:0015850 | organic hydroxy compound transport | | 0.00056 | 0.005189 |
| GO:0007275 | multicellular organism development | | 0.000582 | 0.005354 |
| GO:0032509 | endosome transport via multivesicular body sorting pathway | | 0.000595 | 0.005444 |
| GO:0071985 | multivesicular body sorting pathway | | 0.000595 | 0.005444 |
| GO:0051049 | regulation of transport | | 0.000691 | 0.006272 |
| GO:0019374 | galactolipid metabolic process | | 0.000718 | 0.006446 |
| GO:0019375 | galactolipid biosynthetic process | | 0.000718 | 0.006446 |
| GO:0006997 | nucleus organization | | 0.000718 | 0.006446 |
| GO:0006767 | water-soluble vitamin metabolic process | | 0.000729 | 0.006519 |
| GO:0006397 | mRNA processing | | 0.00074 | 0.006602 |
| GO:0044070 | regulation of anion transport | | 0.000799 | 0.007107 |
| GO:0010274 | hydrotropism | | 0.00083 | 0.007372 |
| GO:0006972 | hyperosmotic response | | 0.000893 | 0.007801 |
| GO:0009738 | abscisic acid-activated signaling pathway | | 0.000904 | 0.007875 |
| GO:0010447 | response to acidic pH | | 0.000928 | 0.008061 |
| GO:0016042 | lipid catabolic process | | 0.000942 | 0.008161 |
| GO:0009668 | plastid membrane organization | | 0.000974 | 0.008358 |
| GO:0010027 | thylakoid membrane organization | | 0.000974 | 0.008358 |
| GO:0006900 | vesicle budding from membrane | | 0.001012 | 0.008661 |
| GO:0044255 | cellular lipid metabolic process | | 0.001042 | 0.00889 |
| GO:0032787 | monocarboxylic acid metabolic process | | 0.001067 | 0.00908 |
| GO:0090630 | activation of GTPase activity | | 0.001129 | 0.009457 |
| GO:0031998 | regulation of fatty acid beta-oxidation | | 0.001129 | 0.009457 |
| GO:0043547 | positive regulation of GTPase activity | | 0.001129 | 0.009457 |
| GO:0046320 | regulation of fatty acid oxidation | | 0.001129 | 0.009457 |
| GO:0051345 | positive regulation of hydrolase activity | | 0.001129 | 0.009457 |
| GO:0006812 | cation transport | | 0.001131 | 0.009457 |
